# Bacterial Stimulation Remodels Macrophage Extracellular Vesicle Lipids and Reveals iNOS as an Inflammatory Cargo

**DOI:** 10.64898/2025.12.11.693607

**Authors:** Miriam Dutkova, Julia Orlovska, Kristyna Turkova, Eva Vojtkova, Marek Oslacky, Lucia Calekova, Veronika Mlcekova, Stepan Strnad, Vladimir Vrkoslav, Vratislav Berka, Josef Chudacek, Stefan Kolcun, Eva Kriegova, Lukas Kubala, Gabriela Ambrozova

## Abstract

**Background:** Inflammation drives the progression of acute and chronic diseases, including sepsis and peritonitis, yet the mechanisms by which bacterial stimuli alter intercellular inflammatory communication remain incompletely understood. Extracellular vesicles (EVs) are increasingly recognized as important mediators of immune signaling and may transmit stimulus-specific lipid and protein cargo between cells. We investigated whether bacterial stimulation remodels macrophage-derived EV composition and whether these changes are associated with altered EV uptake and inflammatory activity.

**Methods:** EVs were isolated from RAW264.7 macrophages under basal conditions or following stimulation with bacterial lysate (BL) from *Lacticaseibacillus rhamnosus* CCM7091 or lipopolysaccharide (LPS). EV identity and quality were assessed according to MISEV recommendations. Lipidomic profiling was performed to define stimulus-associated lipid remodeling, while EV-associated inducible nitric oxide synthase (iNOS) and its enzymatic activity were evaluated using biochemical approaches. EV uptake and functional activity were assessed in macrophages and endothelial cells. EV-associated iNOS was further investigated in murine models of peritoneal inflammation and fibrosis and in EVs isolated from peritoneal exudates of patients with acute peritonitis.

**Results:** Bacterial stimulation altered the molecular composition and biological activity of macrophage-derived EVs. BL-EVs exhibited enrichment of saturated fatty acids and ceramides, generating a distinct lipid signature associated with enhanced cellular uptake. Functionally, bacteria-stimulated EVs promoted macrophage activation, increasing nitric oxide and TNFα production, while LPS-EVs additionally induced endothelial IL-6 and CCL5/RANTES production and ICAM-1 expression. Importantly, enzymatically active iNOS was identified in bacteria-stimulated EVs, demonstrating functional iNOS cargo in small EVs derived from a defined macrophage cell line. EV-associated iNOS was further detected in murine models of peritoneal inflammation and fibrosis and in patient-derived EVs from acute peritonitis, supporting its relevance beyond the *in vitro* system.

**Conclusions:** Bacterial stimulation remodels macrophage-derived EV lipid and protein cargo and generates EV populations with altered uptake and inflammatory activity. These findings identify stimulus-dependent lipid remodeling as a feature of bacteria-stimulated EVs and establish enzymatically active iNOS as a previously unrecognized inflammatory EV cargo. The detection of EV-associated iNOS in experimental models and patients with peritoneal inflammation further supports its potential as a biomarker of inflammatory disease and for therapeutic modulation of EV-mediated immune signaling.

## 1. Introduction

Inflammation is a central driver of pathology in both acute and chronic diseases. Infections such as peritonitis and sepsis are characterized by excessive systemic immune activation, with sepsis alone accounting for an estimated 11 million deaths annually and peritonitis remaining a leading cause of morbidity and mortality in peritoneal dialysis patients. While acute inflammation is indispensable for pathogen clearance, persistent inflammation promotes tissue remodeling and fibrosis, including long-term complications of dialysis and other chronic disorders. Although soluble mediators such as cytokines and chemokines are established contributors, extracellular vesicles (EVs) have recently emerged as additional carriers of pro-inflammatory signals that may exacerbate these processes. Understanding how macrophage-derived EVs are remodeled by bacterial stimulation is therefore of direct clinical relevance.

Macrophages act as sentinels of innate immunity, sensing pathogen-associated molecular patterns (PAMPs) such as lipoteichoic acid (LTA), lipopolysaccharide (LPS), and peptidoglycan via pattern recognition receptors (PRRs). These interactions trigger signaling cascades that initiate both local and systemic inflammation, as observed in sepsis, peritonitis, and inflammatory bowel disease. Beyond cytokine release, activated macrophages upregulate inducible nitric oxide synthase (iNOS), generating nitric oxide (NO), a pleiotropic mediator of antimicrobial defense and immune regulation [1]. Importantly, macrophages also secrete EVs whose molecular cargo reflects their activation state [2]. These vesicles can propagate immune responses both locally and systemically [3, 4], and have been reported to carry cytokines, chemokines, and regulatory RNAs [5–8]. Evidence for iNOS transport by EVs, however, is limited to plasma-derived vesicles [9, 10], with no data available for macrophage-derived EVs.

While the protein and nucleic acid cargo of EVs has been extensively studied, their lipid composition remains comparatively underexplored [11]. Lipids shape vesicle biogenesis, uptake, and stability, and certain species such as ceramides or lysophospholipids directly modulate immune signaling [12–14]. Fatty acid saturation and chain length further determine membrane fluidity and inflammatory potential [15]. In pathological states such as obesity and chronic inflammation, EVs have been implicated in transporting bioactive lipids and free fatty acids [16], yet it remains unknown how infection or PAMP recognition remodels the lipid landscape of macrophage-derived EVs.

Here, we hypothesize that bacterial stimulation reprograms macrophage-derived EVs at the lipid level and examine how these changes affect their uptake and inflammatory activity. We identify saturated fatty acid enrichment and incorporation of iNOS as hallmarks of bacteria-stimulated macrophage-derived EVs, linking bacterial infection to EV remodeling, and demonstrate that iNOS is transported by small EVs not only in *in vitro* systems but also in murine models and clinical samples of peritoneal inflammation. These findings reveal a mechanism by which macrophage-derived EVs amplify systemic inflammation, with implications for sepsis, peritonitis, and other infection-driven pathologies.

## 2. Materials and Methods

We have submitted all relevant data of our experiments to the EV-TRACK knowledgebase (EV-TRACK ID: EV250114) [17].

### 2.1. *Lacticaseibacillus rhamnosus* CCM7091 bacterial lysate (BL) preparation

The BL for the RAW264.7 cell stimulation was prepared as follows: the overnight culture of *L. rhamnosus* CCM7091 (=ATCC 53103), obtained from the Czech Collection of Microorganisms (CCM, Brno, Czech Republic), was inoculated (1% inoculum of culture with OD600nm = 1.0) in de Man, Rogosa and Sharpe broth (MRS) medium (VWR, Czech Republic) and incubated aerobically at 37°C for 48 h. Subsequently, the culture was centrifuged at 8000 x g for 20 min at 4°C, then resuspended in sterile 0.1-µm-filtered phosphate-buffered saline (fPBS; pH 7.5) and centrifuged again (8000 x g, 20 min, 4°C). The pellet was resuspended in sterile deionized H_2_O and incubated at 80°C for 3 h with occasional mixing. After that, the BL was left overnight at −80°C and then heated again at 80°C for 30 min, sonicated for 1 h in water bath and heated again at 80°C for 30 min. Finally, the BL was frozen at −80°C for at least 1h. To check the sterility, the BL samples were incubated both at MRS agar plates and in DMEM, and no bacterial growth was observed. To measure protein concentration in BL samples, Pierce™ BCA Protein Assay (Thermo Fisher Scientific, USA) was performed.

### 2.2. Cell culture

The murine macrophage cell line RAW264.7 (used for isolation of EVs and functional analyses) and murine endothelial cell line MS-1 (used for functional analyses) (American Type Culture Collection, USA) were routinely cultured in DMEM (Gibco, USA), supplemented with 10% (vv^-1^) FBS (Thermo Fisher Scientific, USA), 100 UmL^-1^ of penicillin and 100 μgmL^-1^ of streptomycin (Gibco, USA). Cells were cultivated at 37 °C in a humidified 5% CO_2_ atmosphere.

### 2.3. Isolation of EVs

For isolation of EVs, RAW264.7 cells were seeded in 2 dishes with a surface area of 147.8 cm². After reaching 90% confluency, the cells were harvested into 13 mL of EV-depleted culture medium, prepared by ultracentrifugation (UC) of the whole medium at 175,000 x *g* for 16 h at 4°C (32,000 rpm for rotor SWTi32; Optima TM LE-80K Preparative Ultracentrifuge, Beckman Coulter, USA). Then, they were redistributed evenly into twelve 147.8 cm² dishes, each pre-filled with 24 mL of depleted medium. When the cells reached 70-80% confluence (approximately in 24 h), 6 randomly selected dishes were stimulated with either 100 ngmL^−1^ LPS (*Escherichia coli* O26:B6 LPS, cat. n. 297-473-0, Merck, Germany), or 100 ugmL^-1^ BL, or left untreated to obtain LPS-EVs, BL-EVs or Ctrl-EVs respectively. After 24 h treatment, the media were collected and EVs isolated using differential UC with sucrose cushion purification, modified from [18]. Briefly, conditioned media were collected from six dishes with a surface area of 147.8 cm² and centrifuged at 250 ×⍰g for 5 minutes at 10⍰°C to remove floating cells. Supernatants were transferred to new 50 mL tubes and centrifuged at 1,500 ×1g for 10 minutes at 10 °C to eliminate apoptotic bodies and debris. The resulting supernatants were further cleared at 8,000 ×1g; 4°C for 30 minutes (MR22i Refrigerated Centrifuge, Jouan, USA) for purification from remaining BL and then transferred into new tubes and spun at 14,000 ×⍰g, 10°C for 70 minutes (MR22i Refrigerated Centrifuge, Jouan, USA) to pellet large EVs. Supernatants were poured into 30 mL thin-wall polypropylene ultracentrifuge tubes and balanced using fPBS. UC was performed at 175,000⍰×⍰g for 3 hours at 10⍰°C (32,000 rpm for rotor SWTi32; Optima TM LE-80K Preparative Ultracentrifuge, Beckman Coulter, USA). After removal of the supernatant, pellets were resuspended in 1 mL fPBS and pooled to new conical thin-wall polypropylene ultracentrifuge tubes and diluted with fPBS. A 4 mL layer of sucrose solution (30% sucrose in 20-mM Tris, pH 7.6 in D_2_O, 0.22-µm-filter sterilized) was underlaid to allow removal of protein aggregates in the next spin. Samples were ultracentrifuged at 175,000 ×⍰g for 70 minutes at 10⍰°C with slow deceleration (32,000 rpm for rotor SWTi32; Optima TM LE-80K Preparative Ultracentrifuge, Beckman Coulter, USA). Approximately 5 mL of the sucrose-fPBS interface was collected, transferred to clean ultracentrifuge tubes, diluted with fPBS, and ultracentrifuged again at 175,000 ×⍰g for 16 hours at 10 °C (32,000 rpm for rotor SWTi32; Optima TM LE-80K Preparative Ultracentrifuge, Beckman Coulter, USA). Final pellets of small EVs were re-suspended in ±100 µL of fPBS stored at –80⍰°C in low-binding tubes until further analysis.

The peritoneal EVs from mouse models (peritoneal lavage fluid) and human patient samples (peritonitis exudates) were isolated in a similar way with slight modifications of the first steps of the EVs isolation protocol. First, the floating cells were removed by centrifugation at 250 x g for 5 min at 4°C. The murine samples supernatants from the same treatment group were then pooled (6 mice/treatment for the peritoneal inflammation model; 5 mice/treatment for the peritoneal fibrosis model) and all samples were centrifuged at 1,500 ×⍰g for 10 minutes at 10⍰°C. Supernatants were then refilled up to 30⍰ mL with fPBS and centrifuged at 14,000 ×1g, 10°C for 70 minutes (MR22i Refrigerated Centrifuge, Jouan, USA) to remove large EVs. The supernatants were then transferred to conical 30 mL thin-wall polypropylene ultracentrifuge tubes and the rest of the isolation protocol was carried out as described above.

### 2.4. Visualization of EVs by electron microscopy

4 µL of the EVs sample were applied to freshly plasma-cleaned TEM grids (Quantifoil, Cu, 300mesh, R2/1) and vitrified in liquid ethane using Leica Automatic Plunge Freezer EM GP2 (4°C, 100% rel. humidity, 30-s waiting time, 4-s blotting time and using sensor blotting). The grids were then mounted into the Autogrid cartridges and loaded into Talos Arctica (Thermo Fisher Scientific, USA) transmission electron microscope for imaging. The microscope was operated at 200 kV. The Cryo-TEM micrographs of EVs were collected on the K2 direct electron detection camera at a nominal magnification of 79,000× with an average defocus of −2.2 µm and an overall dose of ∼40 eÅ−2.

### 2.5. Evaluation of size distribution and concentration of EVs by Nanoparticle Tracking Analysis (NTA)

Before analysis, EV samples were diluted 1:50 (EVs from peritoneal exudates 1:10,000) in fPBS by pipetting (10 µL of EVs into 500 µL of fPBS), avoiding vortexing to prevent vesicle aggregation. The diluted samples were then loaded into 1 mL TERUMO Tuberculin (TBC) syringes for measurement. fPBS alone was used as the blank control. The size distribution and concentration of small EVs were analyzed using a NanoSight NS300 Instrument (Malvern Panalytical Ltd, Malvern, UK), equipped with a scientific CMOS (sCMOS) camera and a 488 nm blue laser. Data acquisition and processing were performed using NanoSight NTA software version 3.4 Build 3.4.4. All measurements were carried out using a chamber temperature of 25.0⍰°C with a syringe pump set to a flow rate of 30 µLmin^−1^. For each sample, the camera level was set to 14 and three 60-second videos were recorded under a frame rate of 25 FPS, collecting a total of 1,498 frames. Particle tracking analysis was performed using a detection threshold of 3 and automatic correction for blur size and maximum jump distance. For statistical analysis, ANOVA followed by Dunnett’s multiple comparisons test in GraphPad Prism 8.0.1 (GraphPad Software, San Diego, California, USA) was used; 5-15 independent repetitions were performed.

### 2.6. Determination of EV markers and pro-inflammatory proteins by Western blot (WB)

WB analysis with immunodetection was performed as previously described [19]. For EV samples, EVs were lysed by the addition of a lysis buffer, in a 1:1 ratio (vv^-1^). For MS-1 cell line samples, the frozen cells were lysed in a lysis buffer and sonicated. All the samples were then mixed with Laemmli buffer, heated in a thermoblock at 100°C for 10 min and subjected to SDS-polyacrylamide gel electrophoresis using 10% or 12.5% running gel, based on the detected protein size. For reference, PageRuler Plus Prestained Protein Ladder (cat. n. 26619, Thermo Fisher Scientific, USA) was loaded. The list of antibodies and their dilution is provided in Table S1.

Immunoreactive bands were detected using an ECL™ detection reagent kit (Pierce, USA) by Amersham™ Imager 680 and quantified by scanning densitometry using the Image J™ program. The individual band density value was expressed in arbitrary units and where relevant, the ratio of target protein to housekeeping protein was calculated. Where possible, the values were related to negative control. For statistical analysis, an unpaired t-test to compare two treatment groups or ANOVA followed by Dunnett’s multiple comparison test to compare three groups were used. GraphPad Prism 8.0.1 (GraphPad Software, San Diego, California, USA) was used; 3-8 independent experiments were performed.

### 2.7. Isolation of macrophage membranes for lipidomic analysis

Membrane fractions were isolated from cultured RAW264.7 cells treated or non-treated by BL using a differential centrifugation approach with slight modifications. All procedures were performed on ice to maintain biomolecular stability.

After removing the conditioned medium for EVs isolation, the culture dishes were rinsed with 5 mL of cold PBS to eliminate dead cells and any residual BL, then placed at −801°C for 15 minutes. Subsequently, 3 mL of ice-cold 1x lysis buffer (20 mM HEPES, 1 mM EDTA, 2 mM MgCl_2_, 1 mM DTT, and 250 mM Sucrose) [20] was added to each dish, and cells were left on ice for 10 minutes to swell. Cells were then scraped using a hard cell scraper and all 3 mL were transferred into a pre-chilled Dounce homogenizer. Homogenization was performed with 20 orbital strokes to ensure mechanical disruption.

Lysates from 3 dishes were pooled into a 15⍰mL conical polypropylene tube. The pooled suspension was centrifuged at 1,800⍰×⍰g for 5 minutes at 4⍰°C to pellet nuclei and cellular debris. Supernatants were transferred to Beckman ultracentrifuge tubes, filling each tube to 4.8 mL; an additional 1x lysis buffer was added as needed. Samples were ultracentrifuged at 64,200 x g (23,000⍰rpm for rotor SW55Ti, Optima TM LE-80K Preparative Ultracentrifuge, Beckman Coulter, USA) for 20 minutes at 4⍰°C (acceleration max, deceleration slow). The resulting membrane pellets were gently resuspended in 100⍰µL of 1x lysis buffer and stored at −80⍰°C for analysis.

Protein concentration was determined using Pierce™ BCA Protein Assay (Thermo Fisher Scientific, USA), and samples were adjusted to 11mgmL^-1^ using 1x lysis buffer. The diluted samples were stored at −80°C and lipidomic analysis was performed.

### 2.8. Lipidomic analyses of EVs and macrophage membranes

The EVs (Ctrl-EVs, BL-EVs) and the membranes of their parental cells, either untreated or treated with BL, underwent lipidomic analysis. LC-MS grade solvents and additives were used for sample preparation and analysis: acetonitrile (VWR, PA, USA), water (Fisher Scientific, MA, USA), 2-propanol (Honeywell, NC, USA), formic acid (Thermo Fisher Scientific, MA, USA), ammonium formate (Honeywell, NC, USA), and methanol (VWR, PA, USA).

Lipids were extracted using a modified methyl-tert-butyl ether (MTBE) extraction method [21]. Briefly, samples were placed in 2 mL Eppendorf tubes with 150 μL of cold methanol. Subsequently, 10 μL of EquiSPLASH (Avanti Polar Lipids, AL, USA) was added, and the samples were sonicated for 10 minutes. Next, 500 μL of cold MTBE was added, and the mixture was vortexed at room temperature for 1 hour. Next, 125 μL of water was added to the mixture, which was then vortexed for 5 minutes, and centrifuged for 10 minutes at 500 g. Then, 400 μL of the upper (organic) phase was transferred to a separate vial. The lower phase was re-extracted with 200 μL of a MTBE/methanol/water solvent mixture (10:3:2.5, v/v/v). Extraction step was repeated, and 200 μL of the upper phase was combined with the first upper phase. The combined organic phases were evaporated and reconstituted in 200 μL of methanol/2-propanol (1:1, v/v).

Untargeted lipidomics profiling was performed by coupling a Vanquish UHPLC chromatography system with a TriPlus RTC autosampler to an Orbitrap IQ-X Tribrid mass spectrometer (Thermo Fisher Scientific, MA, USA) equipped with a heated electrospray ionization (HESI). Mobile phase A was acetonitrile/water (60:40 (v/v)), and mobile phase B was 2 propanol/acetonitrile (90:10, v/v). Both mobile phase solutions contained 10 mM ammonium formate and 0.1% formic acid. The separations were performed on the Waters Acquity UPLC BEH C18 column (2.1 x 100 mm, 1.7 μm) operated at 55 °C and at a flow rate of 300 μLmin^-1^. The elution gradient program was as follows: 0 - 2.0 min, 0 - 43% B; 2.0 - 2.1 min, 43 - 55% B; 2.1 - 12.0 min, 55 - 65% B; 12.0 - 18.0 min, 65 - 85% B; 18.0 - 20.0 min, 85 - 100% B; 20.0 - 25.0 min, 100% B; 25.0 - 25.1 min, 100 - 0% B; and 25.1 - 30.0 min, 0% B. The injection volume was 5 μL for positive ion mode and 10 μL for negative ion mode. Full scan and MS/MS (HCD) data were acquired in data-dependent acquisition (DDA) mode separately for each ionization polarity over the m/z range of 250–1600. The MS resolution 120,000 and the MS/MS resolution 15,000 were applied. The electrospray ionization voltage was 3.5 kV, and the capillary temperature was 350⍰°C. Lipid identification was performed using MS-DIAL 5 [22]. Lipidomic data were normalized to protein content measured by BCA assay and to the response of the corresponding internal standards. The known concentrations of the internal standards were used to express the data in absolute concentration units. Finally, the relative abundance of lipid species was calculated as a percentage of the total lipid content for each sample of EVs. The relative representation of fatty acids was calculated as the contribution of each lipid species to the total concentration of the respective fatty acid. Statistical analysis was performed using MetaboAnalyst 6.0 software (http://www.metaboanalyst.ca/). Prior to analysis, the data were subjected to log transformation and Pareto scaling. A volcano plot analysis was used to identify altered lipids (fold change > 2 and p-value < 0.05 by Student’s t-test). The mass spectrometry data have been deposited in the public repository Zenodo: https://doi.org/10.5281/zenodo.17224726.

Lipidomic data were further analyzed using LipidSig 2.0 [23] and Lipid Ontology (LION) enrichment tool to perform differential expression analysis and interpret changes in chain length, degree of unsaturation (double bonds), and to highlight biological/biophysical features of the detected lipids and the potential functional consequences of the detected alterations.

### 2.9. Fluorescent labeling of EVs

EVs were diluted to a concentration of 10^10^ particles per mL in fPBS and labeled by 5-(6) carboxyfluorescein diacetate N-succinimidyl ester (CFSE) (ThermoFisher, cat. n. 65-0850-84). CFSE was added to the EVs in a final concentration of 1mM and incubated for 30 min at 37°C. Subsequently, 3.5 mL of fPBS was added and the suspension was underlaid with 1 mL of 30% sucrose solution (30% sucrose in 20-mM Tris, pH 7.6 in D2O, 0.22-µm-filter sterilized) and ultracentrifuged at 175,000 × g for 2 h at 4°C (38,000 rpm for rotor SW55Ti; slow deceleration; Optima TM LE-80K Preparative Ultracentrifuge, Beckman Coulter; USA). The interface was collected, refilled with fPBS to wash the free label and ultracentrifuged at 175,000 × g for 2 h at 4°C (38,000 rpm for rotor SW55Ti; slow deceleration; Optima TM LE-80K Preparative Ultracentrifuge, Beckman Coulter; USA). The CFSE-labelled MVs were re-suspended in 100 µL of fPBS and immediately used for RAW264.7 treatment. As a negative control, vehicle alone (fPBS) was subjected to the same labelling protocol with CFSE.

### 2.10. Visualization of EV internalization by RAW264.7 cells using confocal microscopy

RAW264.7 cells were seeded at a concentration of 5 x 10^4^ cells per well onto confocal microscopy glass-bottom chambers. The following day, cells were stained overnight with SPY-620-Actin (cat. no. SC402, Spirochrome) (excitation/emission: 619/645 nm) to label the actin cytoskeleton. After incubation, cells were treated with 50 µL of EVs stained with CFSE (excitation/emission: 492/517 nm), corresponding to a dose of 10^5^ particles per cell. Images were acquired 1 h after the addition of EVs, using Olympus IXplore SpinSR SoRa confocal microscope, equipped with dual-channel acquisition (C488/640) for simultaneous detection of CFSE and SPY620 signals. A 60x silica immersion objective was used for imaging. Data were collected under identical acquisition settings and later visualized using Fiji software with Bio-Formats plugin.

### 2.11 Determination of EV uptake using flow cytometry

RAW264.7 cells were seeded at a concentration of 5 x 10^4^ cells per well in a 96-well plate. The following day, cells were treated with 5 µL of CFSE-labeled EVs, corresponding to 10^4^ EVs per cell, and incubated for 1 h at 37°C. After incubation, the medium was removed, cells were washed once with PBS and resuspended in 200 µL of PBS with the addition of 1% FBS. Cell viability was assessed using 1 µL of DAPI (1 µgmL^-1^), added immediately before measurement. Samples were analyzed using BD FACSDiscoverTM S8 Cell Sorter with BD CellViewTM Image Technology. Results are presented as the median of fluorescence intensity (MFI) of viable cells and representative images of positive cells. For statistical analysis, unpaired t-test in GraphPad Prism 8.0.1 (GraphPad Software, San Diego, California, USA) was used; 8 independent repetitions were performed.

### 2.12. Treatment of macrophage and endothelial cells for functional analysis of the EVs

For the functional analyses, either RAW264.7 or MS-1 cells were incubated with the EVs. The RAW264.7 cells were seeded in a 24-well plate in a density of 2.5 × 10^5^ cells per well, whereas the MS-1 cells were seeded in a density of 6.0 × 10^4^ cells per well (24-well plate). 24 h after the seeding, the medium was replaced with a fresh one and the EVs were added to the appropriate wells in a concentration of 2 x 10^3^ particles/cell for RAW264.7 and 1 x 10^4^ particles/cell for MS-1. The 24-hour treatment duration was chosen based on established protocols for measuring macrophage inflammatory responses in our laboratory [24] and allows sufficient time for evaluated protein expression and accumulation of secreted factors. This timepoint captures the robust inflammatory response while maintaining cell viability. As a negative control, the vehicle alone (fPBS) was used; whereas 100 ngmL^-1^ LPS was used for RAW264.7 cells, and 5 ngmL^-1^ IL-1β for MS-1 cells as positive controls. The media were then collected, depleted of any remaining cells (by centrifugation, 300 × g for 5 min at 4°C), and stored at −20°C for further cytokine and RNS detection. The MS-1 cells were washed with ice-cold fPBS and stored at −20°C for WB analysis.

### 2.13. Detection of pro-inflammatory cytokines by ELISA

Levels of the cytokines TNFα, IL-6, and CCL5/RANTES were detected in the cell culture medium collected after the treatment of RAW264.7 or MS-1 cells with the EVs. Detection was performed by ELISA (cat. n. 88-7324-88, 88-7064-88, Thermo Fisher Scientific, USA and MMR00, R&D Systems, Inc., USA) following the manufacturer’s instructions with no modifications. To assess statistical significance, based on the number of compared groups and the character of the data, the following tests were used: TNFα – unpaired t-test with Welch’s correction; IL-6 - Kruskal-Wallis test, followed by Dunn’s multiple comparisons test; RANTES - Ordinary one-way ANOVA with Dunnett’s multiple comparison test. The data were related to negative controls (vehicle-treated cells) and analyzed in GraphPad Prism 8.0.1 (GraphPad Software, San Diego, California, USA); 4-7 independent repetitions were performed.

### 2.14. Measurement of nitrite concentration by Griess reaction

Changes in RNS production were measured indirectly as the accumulation of nitrites, the end product of NO metabolism, in a medium using the Griess spectrophotometric assay, as we described previously [25]. Briefly, after the incubation period, the culture media were collected from the wells and centrifuged at 250 × *g* at 4°C for 5 min. 80 µL of supernatant was then mixed with an equal volume of Griess reagent (Sigma Aldrich, USA) in a 96-well plate and the mixture was incubated at room temperature and in the dark for 30 min. Absorbance was measured at 546 nm (Spectrophotometer Sunrise, Tecan, Switzerland). Sodium nitrite was used as a standard. For statistical analysis, unpaired t-test in GraphPad Prism 8.0.1 (GraphPad Software, San Diego, California, USA) was used; 8-11 independent repetitions were performed and the results are related to the negative control.

### 2.15 iNOS activity assay

The ability of the iNOS enzyme carried by EVs to convert substrate to nitrites was tested using the Nitric Oxide Synthase Activity Assay Kit (Abcam, GB). The assay was performed according to the manufacturer’s instructions with minor alterations to optimize its application for iNOS detection in EVs as follows: 20 μl of NOS Assay Buffer were first added to 20 μl of EVs to disrupt the lipid bilayer and release the encapsulated iNOS enzyme. Next, the samples were incubated according to the manufacturer’s instructions with the enzyme cocktail for 1 hour at 37 °C, NOS Assay Buffer and Enhancer II were added and incubated for 10 min at RT, followed by the addition of the DAN probe. After a further 10-min incubation at RT, Sodium Hydroxide was added to the reaction. Following a final 10-min incubation at RT, fluorescence was measured at Ex/Em = 360/450 nm using a Spark 10M fluorescence spectrophotometer (Tecan, Switzerland). iNOS activity was expressed as relative fluorescence units (RFU). For statistical analysis, Ordinary one-way ANOVA with Dunnett’s multiple comparison test in GraphPad Prism 8.0.1 (GraphPad Software, San Diego, California, USA) was used; 4-5 independent repetitions were performed.

### 2.16 *In vivo* models of peritoneal inflammation and inflammation-connected peritoneal fibrosis

In *in vivo* experiments, male C57BL/6 mice, 10–12 weeks of age and weighing 20–25 g, obtained from the Laboratory of Animal Breeding and Experimental Facility of the Faculty of Medicine (Masaryk University, Brno, Czech Republic) were used. The experiments were approved by the Animal Care Committee of The Czech Academy of Science, Approval No. 94/2019. The mice were maintained in cages with free access to food and drinking water in the certified animal facility with a controlled climate and 12 h/12 h light/dark cycle. For induction of peritoneal inflammation and inflammation-connected fibrosis in peritoneum mice were anesthetized by 2.5%–5% isofluorane Aerrane, the abdomen was shaved and disinfected using 70% ethanol, and pathologic conditions induced as follows. For moderate peritoneal inflammation, purified LPS from *E. coli* O26:B2 in 0.1% BSA in PBS (5mg/kg body weight) and BL from *L. rhamnosus* CCM7091 (16mg/kg body weight) were injected intraperitoneally according to previously published protocols [26]. To induce peritoneal fibrosis, which is associated with inflammation and hypoxia, incisions were made in the skin and peritoneal membrane on the abdominal side of the mice. The intestines were exteriorized, desiccated using surgical gauze pads for 10 minutes, and then gently returned to the abdominal cavity. 100 μL of 0.1 mM CoCl_2_ in PBS was applied into the cavity, and the abdominal incision was closed [27]. Local anesthetic cream Emla (Astra Zeneca, UK) was spread over the abdomen to ease the pain.

After induction, mice were maintained for 2 (peritoneal inflammation) or 3 days (peritoneal fibrosis), after which they were anesthetized, and peritoneal lavage fluid was obtained by washing the peritoneal cavity with 4 mL of cold fPBS. Peritoneal lavages were centrifuged at 250 x g for 10 min and the pelleted cells were analyzed using clinical hemocytometer (Mindray, China). The supernatants were subjected to peritoneal EVs isolation as described in chapter 2.3. Mice were sacrificed by cardiac puncture; blood was collected into EDTA tubes for the detection of leukocyte count. In the peritoneal fibrosis model, representative mice were maintained for 10 days post-induction. After sacrifice, fibrotic tissue along with surrounding structures was excised from the peritoneal cavity and fixed in 4% paraformaldehyde, stained with hematoxylin and eosin and Masson’s trichrome for collagen detection according to standard protocols. Immunohistochemistry was performed using α-Smooth Muscle Actin antibody (Cell Signaling) and secondary anti-rabbit antibody conjugated with Alexa Fluor 555 (Abcam), Roti-Mount FluorCare DAPI (Carl Roth) and scanned using a Leica TCS SP8X confocal microscope (Leica Microsystems).

For statistical analysis of the hemocytometric analysis, ANOVA followed by Bonferroni’s multiple comparisons test in GraphPad Prism 8.0.1 (GraphPad Software, San Diego, California, USA) was used; 5-6 independent repetitions were performed.

### 2.17 Clinical peritonitis study cohort

In this study, a pilot subcohort of four adult patients with surgically confirmed acute secondary peritonitis (type diffuse purulent peritonitis) requiring emergency operative source control was included. Intraoperative peritoneal swabs were collected from the most inflamed parietal and visceral peritoneal surfaces, including exudate when present, and processed for microbiological analysis and EVs isolation as described in chapter 2.3. Clinical data included inflammatory markers, comorbidities, type of infection, treatment, operative findings and 90-day outcome.

All procedures performed in this study were conducted in accordance with the Declaration of Helsinki. Informed consent was obtained from all individual participants included in the study. The study protocol was approved by the Ethics Committee of University Hospital Olomouc and Palacky University Olomouc [ID 100/26].

### 2.18 Statistical analysis

Data are presented as mean ± SD. All measurements were performed at least in duplicate in *n* independent experiments. The statistical significance was assessed at \**p* < 0.05. Details regarding statistical analyses can be found in the description of the methods.

### 2.19 AI tools

Artificial intelligence (AI) tools were used solely to assist with textual editing and language refinement in this manuscript. All experimental design, data acquisition, analysis, and interpretation, as well as the figures and original scientific content, were independently performed by the authors and represent original work.

## 3. Results and discussion

### 3.1. Effects of bacterial stimuli on EV production and morphology

While macrophages are well known for their inflammatory responses, the effects of bacterial stimulation on their EV release and physical properties remain unclear. Therefore, we isolated EVs from macrophage cell line RAW264.7 treated with two bacterial stimuli: LPS, a standard pro-inflammatory component of the G-bacterial cell wall [28–31] and BL from the G+ bacterium *L. rhamnosus,* which provides a much more complex mixture of microbial-associated molecular patterns than purified LPS and may therefore provide more comprehensive stimulation [32]. After characterizing the BL composition (Fig. S1) and confirming the macrophage activation following LPS and BL stimulation (Fig. S2), we isolated EVs at 24 h from treated and untreated cells (LPS-EVs, BL-EVs, and Ctrl-EVs). We characterized the EVs following the MISEV2023 guidelines [33]: measured size distribution and concentration with NTA (Fig. 1a-d), visualized the extremities (Ctrl-EVs and LPS-EVs) by Cryo-EM (Fig 1e) and detected typical EV markers (including negative EV markers) by WB (Fig. 2a,b). The average concentration was about 8 to 9×10^10^ particles/mL, corresponding to 120-145 particles per frame, and the mean particle size ranged from 110-125 nm, consistent with the Cryo-EM pictures (Fig. 1e). Full results from NTA measurements for all three EVs groups are provided in Table S2. Although reported EV concentrations vary between studies due to differences in starting volume, our size range aligns with prior EV reports [28, 34, 35]. To ensure proper standardization, EVs counts were normalized to the number of live maternal cells per mL (LML). After this adjustment, no significant difference in particle number were detected between the treatment groups (Fig. 1c), although we cannot rule out the possibility of subtle changes below the detection threshold. Similarly, EVs released from LPS- and BL-treated cells showed a trend toward reduced size; however, this decrease did not reach statistical significance. The size trend accords with Emmerson et al. 2024, who observed smaller RAW264.7-EVs after *Salmonella* Typhimurium infection [36], and with reports of reduced size after M1-like polarization [37]. Although LPS has been reported to increase EV concentration *in vivo* in bronchoalveolar lavage fluid [38], the *in vitro* findings are heterogeneous. Jin et al. 2023 reported time-dependent changes in LPS-EVs release, with increased production after 8 h but decreased after 24 h compared with controls [39], suggesting that overall EV levels remain relatively in line with our observations. In contrast, LPS-treated BMDM released higher EV concentration at 6, 12, and 24 h [40]. Another study using RAW264.7 cells indicated a possible reduction in LPS-EVs production; however, this conclusion was based mainly on reduced mRNA expression of EV biogenesis markers, while the actual EV concentrations were not statistically different [28]. Taken together, these data show that *in vitro* results on LPS-EV release are inconsistent and likely influenced by cell type and time. Given this relative stability in EV production, we next examined whether bacterial stimulation alters the molecular cargo of EVs.

**Figure 1.**
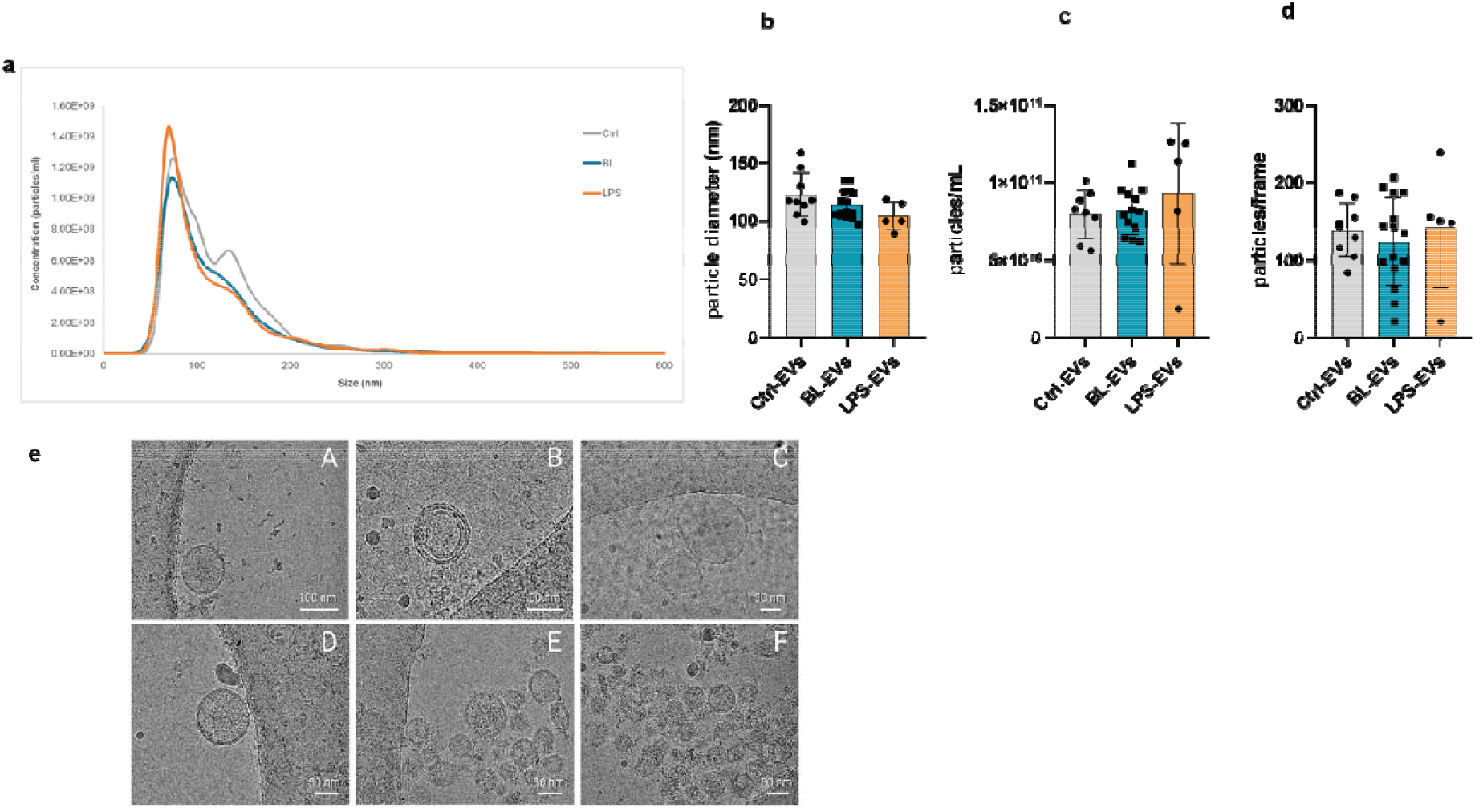
Biophysical characterization of macrophage-derived extracellular vesicles (EVs) **(a-d) Particle size and concentration of EVs measured by Nanoparticle Tracking Analysis (NTA).** EVs isolated from bacterial lysate-treated (BL-), lipopolysaccharide-treated (LPS-) or untreated control (Ctrl-) macrophages were analyzed. Particle concentration was normalized to live maternal cells per mL (LML). Data are shown as mean ± SD. For statistical analysis, Ordinary One-way ANOVA followed by Dunnett’s multiple comparisons test versus Ctrl-EVs was performed. \**p* < 0.05; n = 5-15. **(e) Cryo-Electron microscopy of EVs.** Representative images of Ctrl-EVs (A-C) and LPS-EVs (D-F).

**Figure 2.**
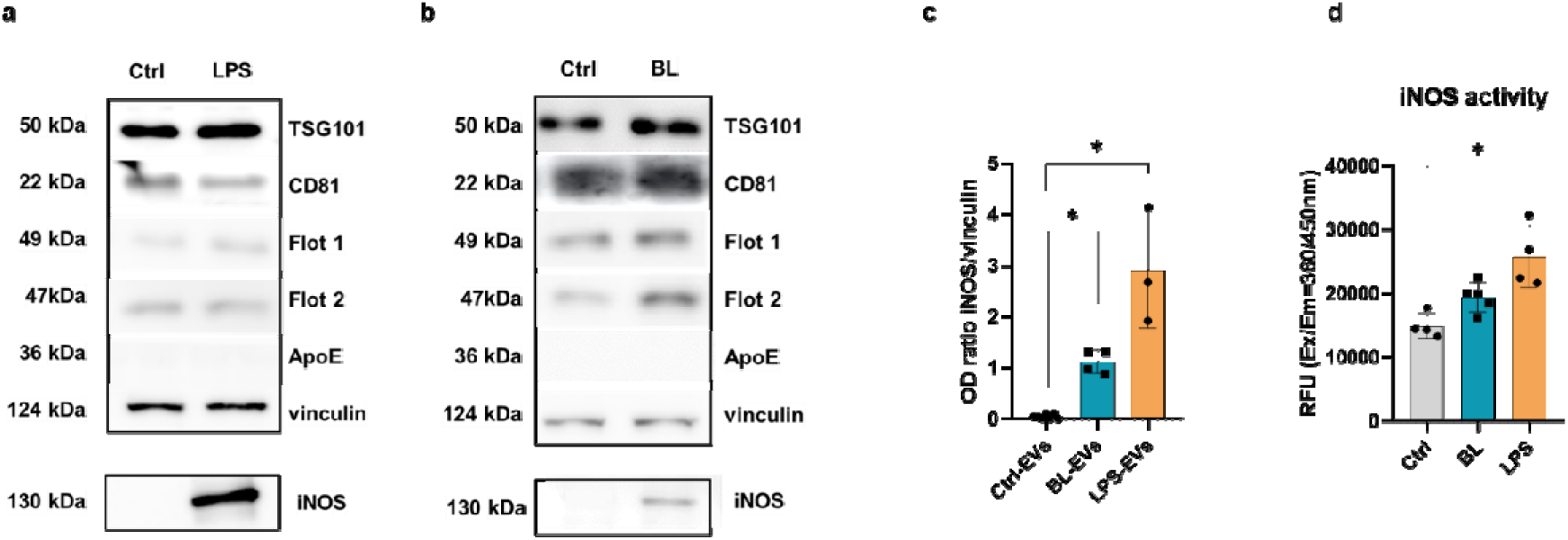
Characterization of macrophage-derived extracellular vesicle (EV) cargo. **(a-b) Detection of EV markers by Western blotting.** EVs isolated from bacterial lysate-treated (BL-), lipopolysaccharide-treated (LPS-) or untreated controls (Ctrl-) macrophage-derived EVs were analyzed. The expression of EV markers, including negative EV marker was detected in BL-EVs, LPS-EVs and Ctrl-EVs. Representative blots are shown. **(c) Relative abundance of iNOS in EVs.** Ratio of optical density (OD) for iNOS/vinculin was quantified. Data are presented as mean ± SD. Ordinary one-way ANOVA followed by Dunnett’s multiple comparison test versus Ctrl-EVs was performed. \**p* < 0.05; n = 3-7. **(d) EV-associated iNOS retains enzymatic activity following macrophage activation.** iNOS enzymatic activity was measured in EVs derived from untreated control (Ctrl), BL-stimulated, and LPS-stimulated macrophages and is presented as relative fluorescence units (RFU; Ex/Em 360/450 nm). Data are presented as mean ± SD. Ordinary one-way ANOVA followed by Dunnett’s multiple comparison test versus Ctrl-EVs was performed; \**p* < 0.05; n = 4-5.

### 3.2. Bacterial stimulation remodels EV protein cargo and enriches EVs with enzymatically active iNOS

Beyond confirming the presence of standard EV markers (TSG101, CD81) and the absence of ApoE [33], we also detected vinculin (Fig. 2a,b) a membrane-associated cytoskeletal protein that anchors actin filaments to transmembrane receptors such as integrins, thereby contributing to the structural organization and stability of the EV membrane [41]. Moreover, we identified additional cargo changes. We detected flotillin-1 and flotillin-2, with flotillin-2 being selectively increased in EVs from LPS- and BL-treated macrophages (Fig. 2a,b) but non-changed in parental cells (Fig. S3). To our knowledge, this selective enrichment has not been previously reported. Such changes may reflect altered EV biogenesis during macrophage activation, with potential effects on cargo sorting and signaling. Enhanced flotillin-2 could contribute to the pro-inflammatory activity of these EVs, possibly through lipid regulation and TLR signaling [42–44], although further studies are required.

In line with this pro-inflammatory signature, we next examined the presence of iNOS in EVs. Interestingly, iNOS levels were significantly increased in EV preparations from LPS- and BL-stimulated macrophages, compared with the controls (Fig. 2c). Earlier studies by Azevedo et al. 2007 [45] first reported iNOS in EV-rich plasma and platelet fractions, and more recently Webber et al. 2019 [9] confirmed the presence of iNOS in EV-containing samples from blood. These findings, obtained in complex *in vivo* sources, suggested that iNOS can associate with EVs. To our knowledge, however, iNOS has not previously been identified in small EVs and EVs derived from a defined cell line. Our data therefore extend these earlier observations to a controlled *in vitro* model, reinforcing the physiological relevance of iNOS packaging into EVs.

To determine whether EV-associated iNOS retains its enzymatic function, we next assessed its activity. EVs derived from BL-stimulated macrophages showed only a modest increase in iNOS activity. Notably, EVs isolated from LPS-stimulated macrophages exhibited significantly higher iNOS activity than the negative control (Fig. 2d), demonstrating that EV-associated iNOS remains enzymatically active. To our knowledge, this is the first report of enzymatically active iNOS associated with small EVs. This finding raises the possibility that EVs may serve not only as carriers of iNOS protein but also as vehicles for its catalytic activity, potentially facilitating NO production and propagating inflammatory signaling beyond the site of macrophage activation. Future studies will be required to determine whether EV-associated iNOS produces biologically relevant levels of NO in recipient cells and contributes to downstream inflammatory responses.

### 3.3. Lipidomic profiling reveals distinct remodeling of macrophage membranes and EVs upon bacterial stimulation

Because the EV lipid membrane is a structural determinant of vesicle size, cargo loading, and interactions with recipient cells, we next performed comprehensive lipidomic profiling of both maternal macrophage membranes and their derived EVs. To our knowledge, studies systematically addressing EV lipid composition under inflammatory or bacterial conditions remain scarce [12, 36]. Whereas the impact of LPS on macrophage and EV lipid profiles has been described [28, 46], the effect of more complex G+ stimuli such as BL from *L. rhamnosus* has not been investigated. We therefore compared the lipid profiles of BL-treated (MBL) versus control (MCtrl) macrophage membranes, and of their EVs (BL-EVs, Ctrl-EVs) (Fig. S4).

In the maternal macrophage dataset, the PCA, lipid heatmaps and volcano plot demonstrated a clear separation between MBL and MCtrl profiles (Fig. S5). BL exposure induced a global upregulation of multiple lipid classes, including phosphatidylglycerols (PGs), phosphatidylinositols (PIs), PCs, lysophosphatidylcholines (LPCs), and ceramides, suggesting a stress-activated, pro-inflammatory lipid metabolism. However, some of the effects in membranes can be addressed to bacterial contamination. Such as the increase in PGs, one of the most abundant lipid classes with phosphatidylethanolamine (PE) and cardiolipin (CL) in bacteria (Fig. S6a) [47–49]. CLs are enriched in short and saturated lipid species which can be also attributed to contamination but there is not a clear pattern in PEs and no digalactosyldiglyceroles (DGDG) were identified. This indicates that not only contamination is the factor differentiating BL and controls but lipid remodeling because of pro-inflammatory macrophage reaction. The most obvious is the increase in lysophospholipids (LPC, LPE, LPA) which act as signaling molecules during inflammation [50, 51]. The LPCs role in inflammation is complex and can have pro-inflammatory as well as anti-inflammatory effects. The pro-inflammatory effect is attributed to saturated and monounsaturated LPC (LPC 16:0, LPC 18:0, LPC 18:1) while the latter one to PUFAs containing LPCs (LPC 22:4 etc.)[52]. The LPC 16:0 and LPC 18:0 are strongly enriched in BL treated membranes indicating pro-inflammatory reaction (Fig. S6b). The LPC activates the TLR4 [53] and have interacts with diverse cell types including macrophages as well as other immune cells [54]. TLR4 can be activated by saturated FAs [55, 56]. The observed increase in ceramides can be seen as a pro-inflammatory reaction because they are potent pro-inflammatory agents [57]. The sphingolipid metabolism has been shown to be activated via TLR4 and as a result increased ceramide concentration was observed [56]. In contrast, control cells exhibited consistently lower levels of these lipid classes, indicating their quiescent state.

LION (Lipid Ontology) enrichment analysis was applied to functionally interpret the lipidomic data by linking lipid species to ontology terms describing classification, biophysical properties, subcellular localization, and biological function. Analysis confirmed an enrichment in ceramides and signaling lipids in MBL, accompanied by remodeling of glycerophospholipids and lysophospholipids (Fig. S7). This lipid reprogramming suggests increased membrane-associated signaling activity and stress responses, including pro-inflammatory action, together with structural changes that could influence membrane rigidity, curvature, and downstream interactions with the extracellular environment.

Also in EVs, PCA and heatmap revealed a strong separation between BL-EVs and Ctrl-EVs (Component 1: 60.4% variance explained) (Fig. 3a,b). BL-EVs were enriched in ceramides (e.g., Cer 18:1;O2/16:0, Cer 18:1;O2/24:0), PIs (PI 16:0_18:1, PI 18:0_18:1), and PCs (PC 14:0_16:1) (Fig. 3a-c). Ceramides are well-recognized mediators of inflammation, apoptosis, and M1-like polarization [58], while altered PI and PE metabolism has been linked to EV biogenesis and signaling [13]. Interestingly, unlike in other inflammatory models [13], we did not observe PS upregulation in BL-EVs, suggesting selective remodeling rather than a generalized apoptotic signature.

**Figure 3.**
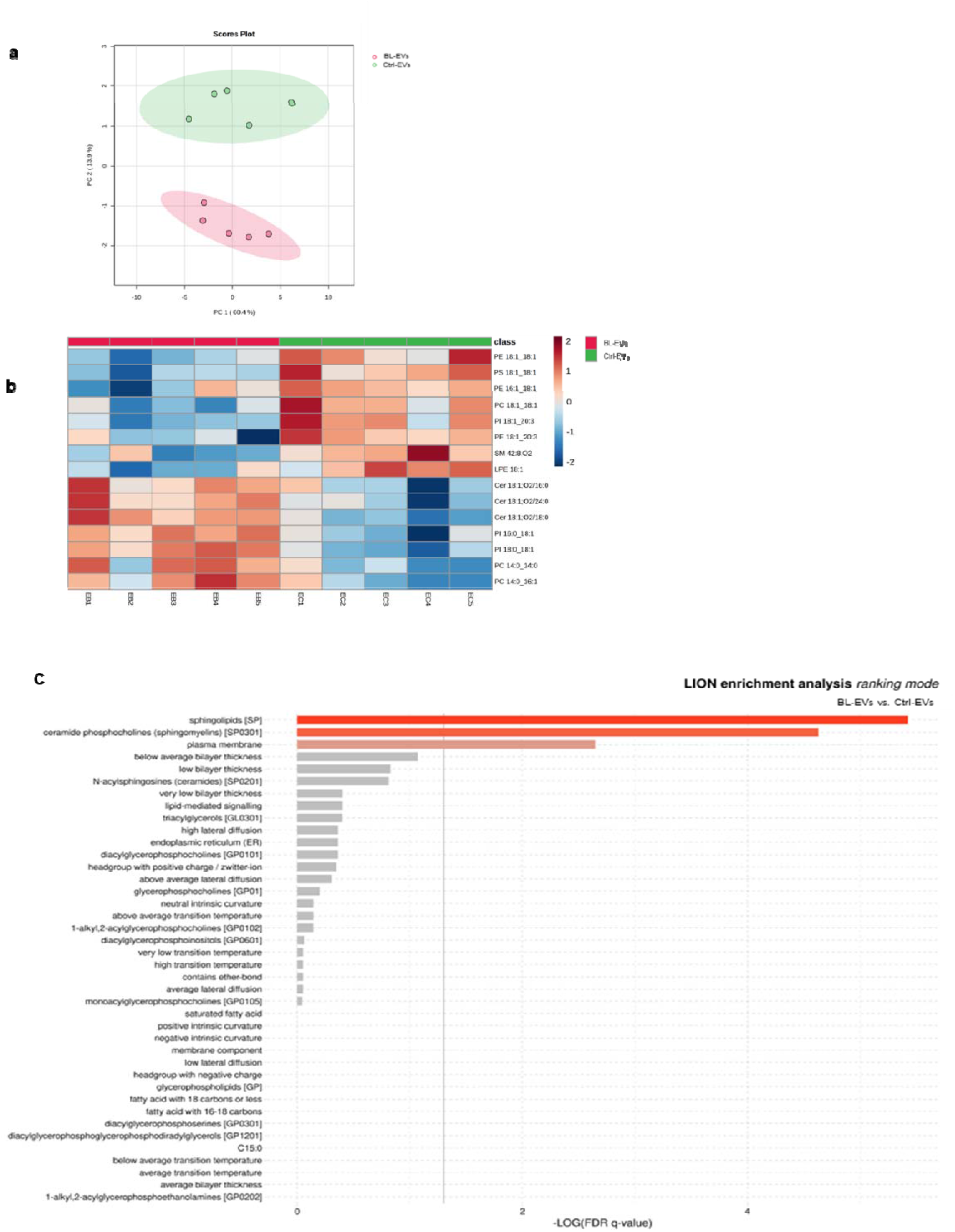
Lipidomic profiling of macrophage-derived extracellular vesicles (EVs). **(a) Principal Component Analysis (PCA) of lipidomic profiles comparing EVs isolated from bacterial lysate-treated (BL-) and untreated control (Ctrl-) macrophages**. Each point represents one biological replicate (n = 5). **(b) Heatmap of the top 15 differentially abundant lipid species in BL-EVs versus Ctrl-EVs**. Data are shown as log₂ fold change. **(c) Lipid Ontology (LION) enrichment analysis comparing BL-EVs and Ctrl-EVs, highlighting biological and biophysical features associated with altered lipid composition**. Enrichment significance was determined using adjusted p-values (FDR < 0.05).

Importantly, Ctrl-EVs retained a composition enriched in LPE, and PE species indicative of a homeostatic lipid balance. This provides a clear baseline against which BL-induced remodeling can be evaluated. Although studies specifically addressing lipidome composition of EVs are limited [12], a recent study by Emerson et al. 2024 performed lipidomics on EVs from RAW264.7 cells following *Salmonella* infection [36]. They observed infection-induced alterations in sphingolipids (e.g., ceramides, sphingomyelin) and glycerophospholipids (e.g., PCs, PIs), accompanied by reduced vesicle size. Notably, similar to our results, most lipids enriched in BL-EVs were already present in Ctrl-EVs, suggesting that bacterial stimulation primarily reshapes the abundance of existing lipid classes rather than introducing new species. This consistency across pathogens reinforces our interpretation that lipid remodeling is quantitative rather than qualitative, yet sufficient to alter EV properties. Concerning potential contamination of BL-EVs by BL, these EVs lacked PGs (Fig. 3b), one of the most abundant lipid classes in bacteria and a potential bacterial contaminant identified in the membranes of BL-treated maternal cells, which indicates that BL-EV preparations were free of contamination.

LION analysis of EVs further demonstrated enrichment in sphingolipids, sphingomyelins, and ceramides, together with plasma membrane-associated lipid signatures (Fig. 3c). Such remodeling implies increased membrane rigidity and raft-like organization, which could enhance EV stability while modulating uptake and signaling in recipient cells.

Together, these results demonstrate that bacterial stimulation drives broad lipid remodeling in both macrophage membranes and their EVs, characterized particularly by shifts in glycerophospholipids, sphingolipids, and ceramides. Because FAs are the structural units underlying these lipid classes and strongly determine membrane fluidity, curvature, and signaling potential, we next focused specifically on FA composition as a critical dimension of this remodeling.

### 3.4. Bacterial stimulation alters fatty acid composition within lipidomic remodeling of macrophage membranes and EVs

As a focused extension of the lipidomic dataset, we examined FA composition in macrophage membranes and EVs. FAs are integral components of glycerophospholipids and sphingolipids, and their saturation level is a major determinant of membrane fluidity, rigidity, and downstream signaling. We hypothesized that BL-induced remodeling of lipid classes would be accompanied by enrichment in short-chain and saturated FAs, consistent with the pro-inflammatory profile inferred from LION analysis.

Primary screening of changes of carbon chain length (total C index) and unsaturation level (total DB index) of macrophage membranes and EVs revealed non-significantly diminished double bonds counts in both membranes and EVs from BL-stimulated cells when compared to control ones and reduction in carbon chain length in MBL membranes (Fig. S8a, b; Fig. 4a, b). Further, distinct clustering of BL-treated versus Ctrl samples was observed in both macrophage membranes and EVs by PLS-DA (Fig. S8c, Fig. 4c). In macrophage membranes, BL exposure increased the abundance of both very-long-chain (e.g., C20:0, C26:0) and shorter saturated FAs (e.g., C10:0, C8:0), accompanied by a trend toward reduced double bond and carbon indices, although these changes did not reach statistical significance. This remodeling indicates a shift toward less fluid, more rigid membranes, a condition classically linked to impaired receptor mobility and the promotion of pro-inflammatory lipid microdomains [59, 60]. Although the mechanistic link between FA remodeling and inflammatory signaling remains incompletely understood, our data are consistent with prior reports showing that increase of saturated FA (reduced double bonds) enhances pro-inflammatory macrophage activation [61]. The enrichment of membrane component lipids further points to changes in lipid packing and curvature that could facilitate EV budding or other membrane-driven processes during macrophage activation [62, 63].

**Figure 4.**
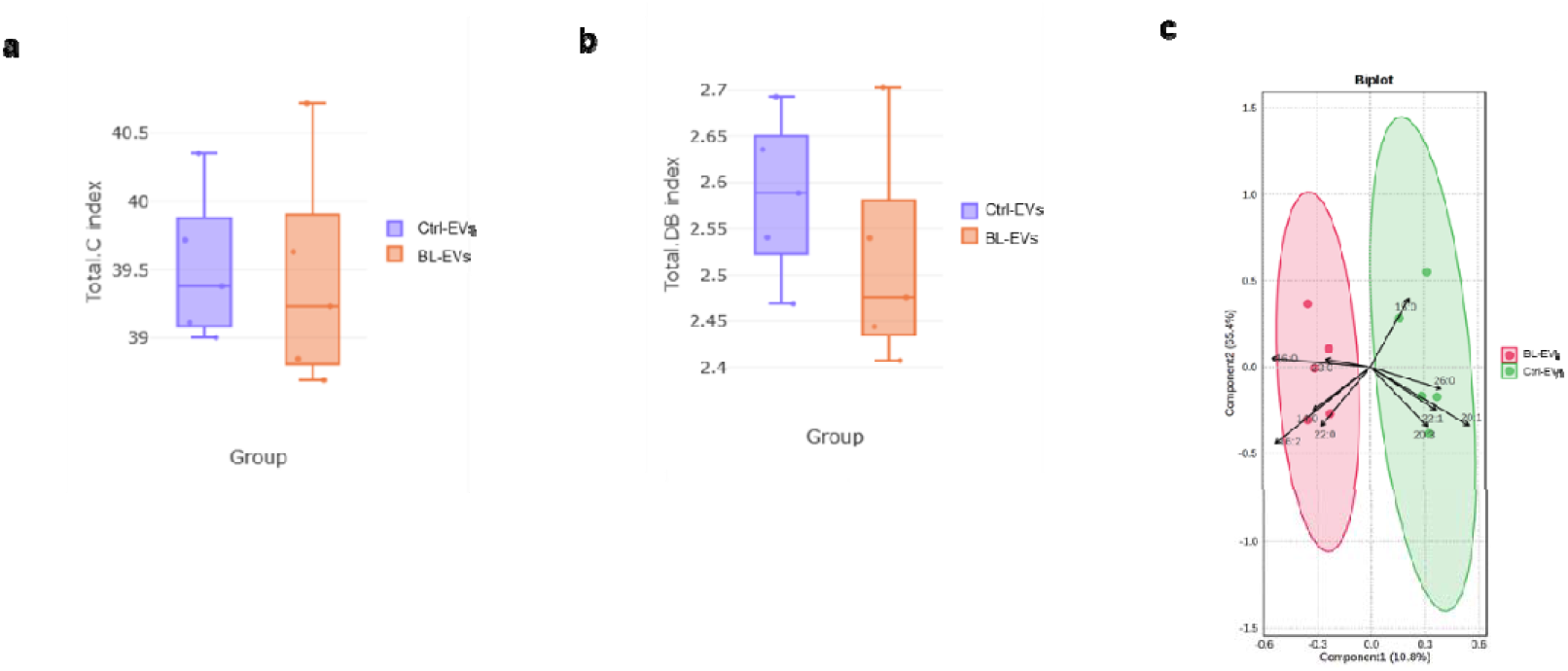
Fatty acid (FA) composition of macrophage-derived extracellular vesicles (EVs). **(a) Differential expression analysis of FA carbon chain length in EVs isolated from bacterial lysate-treated (BL-) and untreated control (Ctrl-) macrophages, performed using LipidSig 2.0.** **(b) Differential expression analysis of FA degree of unsaturation in BL-EVs versus Ctrl-EVs, performed using LipidSig 2.0.** **(c) Biplot of Partial Least Squares Discriminant Analysis (PLS-DA) showing separation of FA composition between BL-EVs and Ctrl-EVs samples.** Each point represents one biological replicate (n = 5).

In EVs, BL similarly enriched saturated FAs (e.g., C23:0, C22:0, C16:0, C14:0), with only a minor reduction in unsaturation (e.g. 16:2) (Fig. 4a-c). Lower double bond content decreases membrane fluidity and increases lipid order, which may enhance EV stability and potentially facilitate fusion or uptake by recipient cells [64]. Additionally, the enriched saturated lipid content may contribute to the pro-inflammatory potential of these BL-EVs either by directly engaging PRRs such as TLRs on target cells or by altering membrane microdomains to prime signaling pathways [65, 66]. This lipid remodeling likely affects both the biophysical properties and biological function of the BL-EVs, thereby enhancing their pro-inflammatory capacity in the context of observed transport of pro-inflammatory cargo (iNOS) (Fig. 2). Thus, these compositionally altered BL-EVs are not only reflective of a pro-inflammatory state in the donor macrophages but may also act as active mediators of inflammation by efficiently delivering pro-inflammatory signals to surrounding or distant cells.

Taken together, FA remodeling toward more saturated species reinforces the broader lipidomic evidence of ceramide- and sphingolipid-driven membrane rigidification which may enhance EV stability and uptake while amplifying their pro-inflammatory potential. While macrophage membranes displayed some signatures suggestive of bacterial contamination, the EV fraction appeared largely free of such components, as none of the previously noted bacterial lipids were detected. Thus, BL-EVs reflect genuine macrophage-derived changes that reduce membrane fluidity, increase lipid order, and enhance pro-inflammatory signaling. Future studies should directly test how ceramide and FA enrichment affect EV uptake, stability, and immune activation.

### 3.5. BL-EVs are internalized more efficiently than Ctrl-EVs

To test whether the changes in EV lipid composition after bacterial stimuli could enhance fusion and uptake by recipient cells, we assessed EV internalization. Although CFSE labeling efficiency was comparable between Ctrl-EVs and BL-EVs (data not shown), when CFSE-labeled Ctrl- and BL-EVs were added to RAW264.7 macrophages at equal particle numbers (10⁵ particles per cell), substantially greater uptake of BL-EVs was observed by confocal microscopy (Fig. 5a). Flow cytometry confirmed this finding, showing a significant increase in BL-EVs internalization compared with controls (Fig. 5b,c). These results suggest that bacterial stimulation remodels EV composition and properties in a manner that facilitates uptake and strengthens intercellular communication.

**Figure 5.**
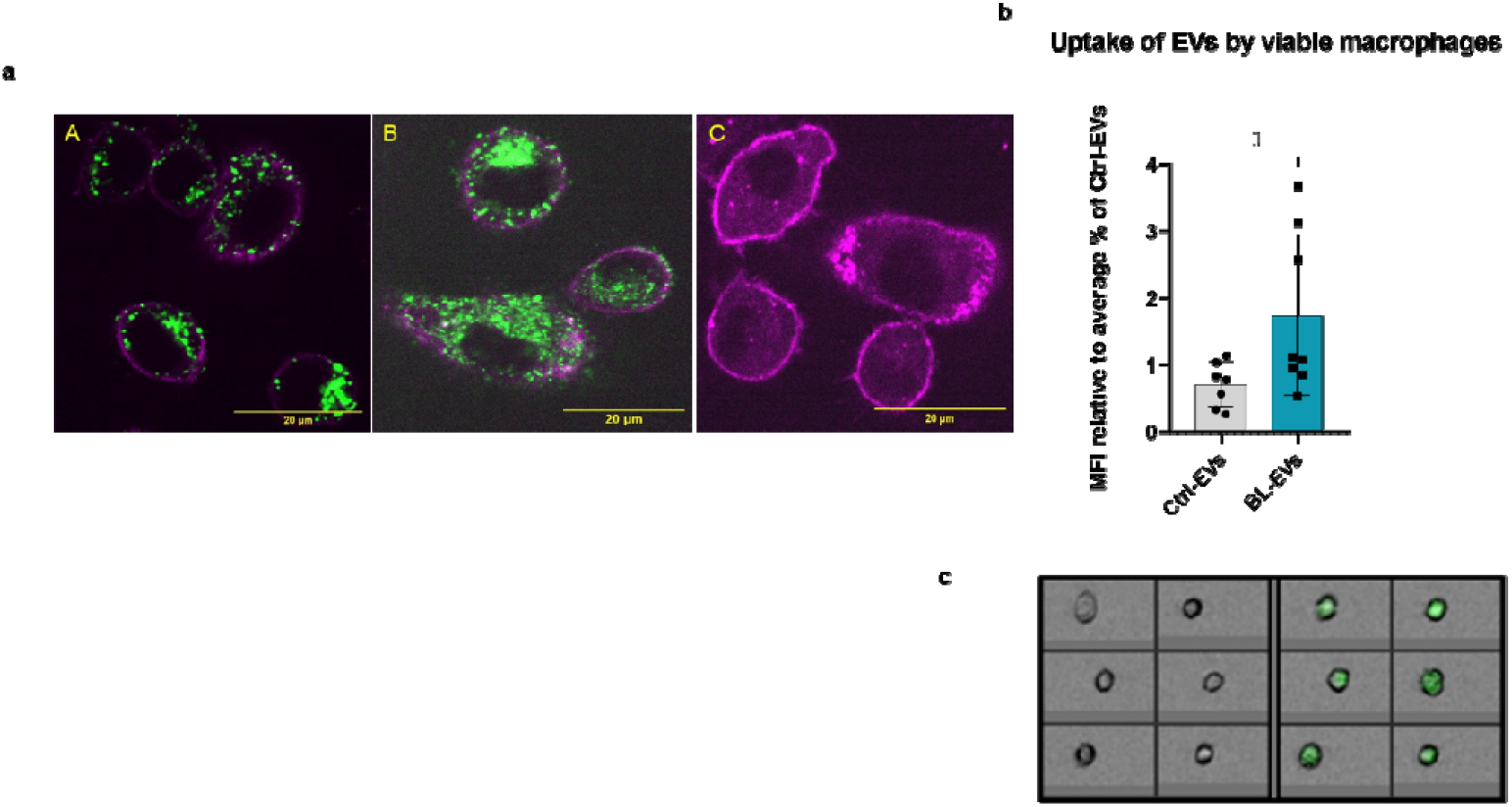
Interaction of macrophage-derived extracellular vesicles (EVs) with target macrophages. **(a) Visualization of EV uptake by macrophages using confocal microscopy**. RAW264.7 cells were incubated with CFSE-labeled Ctrl- or BL-EVs. The cytoskeleton was stained with SPY-620-Actin (pink), and EVs were visualized by CFSE (green). A: Ctrl-EVs, B: BL-EVs, C: PBS control (subjected to identical labeling protocol). **(b-c) Quantification of EV uptake by macrophages**. Flow cytometry analysis of CFSE-labeled Ctrl- and BL-EVs incubated with RAW264.7 cells. Uptake was expressed as median fluorescence intensity (MFI) of viable cells, normalized to average percentage of Ctrl-EVs. Representative images of CFSE-positive (Ctrl-EVs-treated) and CFSE-negative (PBS-treated) cells are shown. Cell viability was assessed using DAPI (1ug/mL). Data are presented as mean ± SD. For statistical analysis, unpaired t-test was performed. \**p* < 0.05; n = 8.

Consistent with this observation, endothelial cell-derived EVs released after LPS stimulation are also taken up more efficiently than controls [67]. Mechanistically, the lipid composition of EV membranes can strongly influence uptake. More rigid membranes may favor fusion, whereas EVs enriched in saturated lipids and thus more rigid may preferentially enter via slower endocytic routes or be cleared through phagocytosis – a pathway particularly relevant in macrophages [68, 69], thereby increasing their potential impact on downstream immune signaling. Together with previous evidence that bacterial exposure reshapes EV biogenesis and cargo [70, 71], our data indicates that bacterial infection-driven remodeling of EVs not only modifies their molecular content but also enhances their capacity for cellular uptake.

### 3.6. BL-EVs amplify pro-inflammatory signaling in recipient macrophages

After demonstrating that BL-EVs are internalized more efficiently than Ctrl-EVs, we next asked whether this enhanced uptake translates into functional consequences. When RAW264.7 macrophages were exposed to Ctrl- or BL-EVs for 24 h, we observed significant upregulation of classical inflammatory markers. Specifically, BL-EV treatment increased RNS production, as is evident from upregulated iNOS protein expression (Fig. 6a) and enhanced nitrites levels (Fig. 6b). Moreover, TNFα secretion was significantly elevated in BL-EV–treated macrophages compared with controls (Fig. 6c). These results parallel the known effects of LPS-EVs [28, 30, 72], supporting the idea that macrophage-derived EVs produced under bacterial stimulation act as amplifiers of inflammation and may propagate the immune response beyond the directly infected cells.

**Figure 6.**
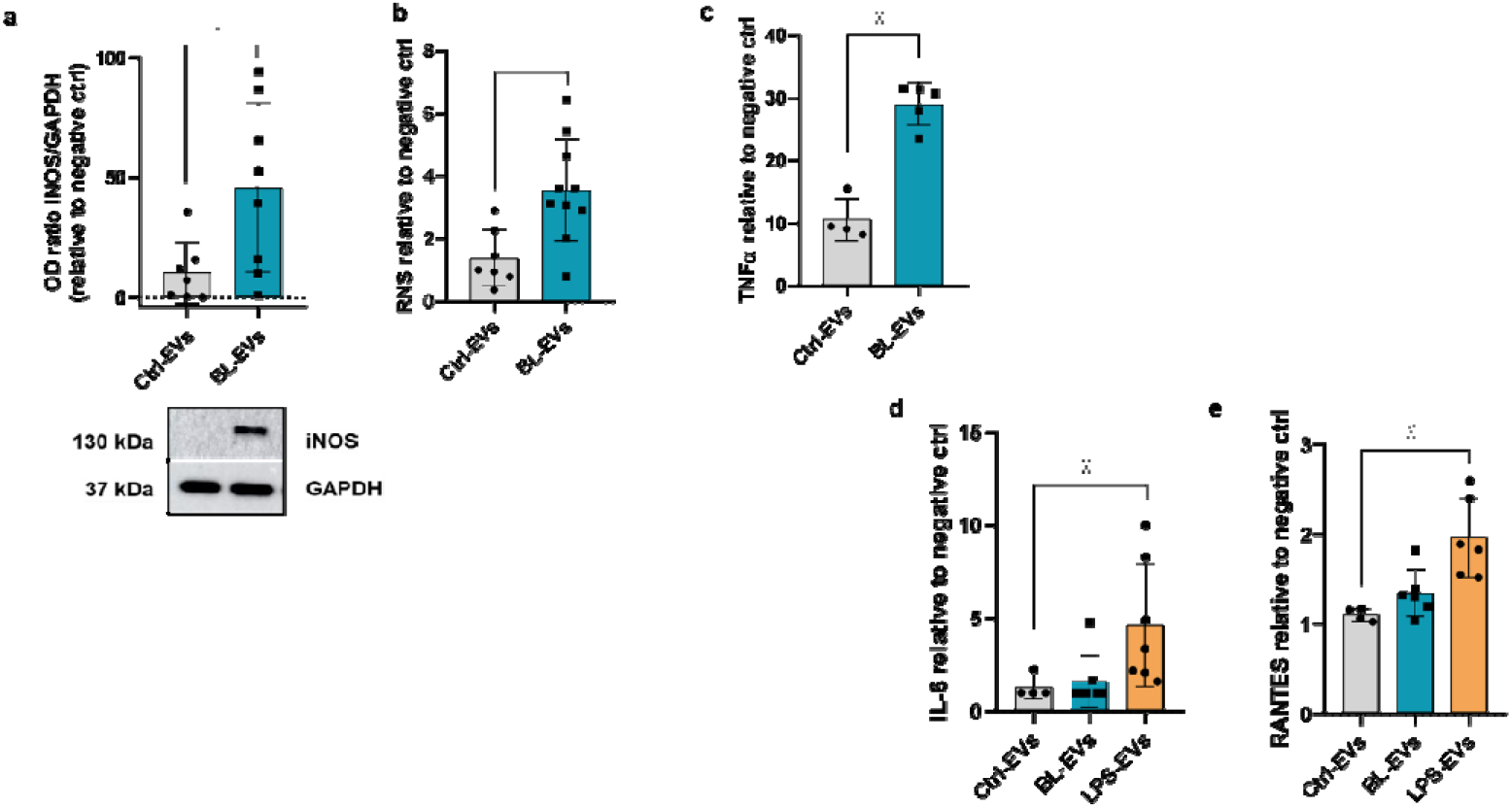
Interaction of macrophage-derived extracellular vesicles (EVs) with target macrophages and endothelial cells. **(a) Induction of iNOS in macrophages treated with EVs**. RAW264.7 cells were treated with Ctrl- or BL-EVs for 24 h, and iNOS expression was analyzed by Western blotting. Representative blots for iNOS and GAPDH are shown. Densitometry ratios (iNOS/GAPDH) were normalized to negative control (vehicle-treated cells, fPBS). Data are presented as mean ± SD. For statistical analysis, unpaired t-test was performed. \**p* < 0.05; n = 8. **(b-c) Production of reactive species of nitrogen (RNS) and TNFα in macrophages treated with EVs**. RAW264.7 cells were treated with Ctrl- or BL-EVs for 24 h. RNS levels in media were measured by Griess reaction and TNFα by ELISA. Values were normalized to vehicle control (fPBS). Data are presented as mean ± SD. For statistical analysis, unpaired t-test was performed for the RNS data, and unpaired t-test with Welch’s correction for the TNFα data. \**p* < 0.05; n = 8-11 for RNS, n = 4-5 for TNFa. **(d-e) Endothelial activation by EVs**. MS-1 endothelial cell line was treated with Ctrl-, BL-, or LPS-EVs for 24 h. IL-6 and RANTES levels in supernatants were quantified by ELISA and normalized to vehicle control (fPBS). Data are presented as mean ± SD. For statistical analysis of the IL-6 data, Kruskal-Wallis test followed by Dunn’s multiple comparisons test was performed; for the RANTES data, Ordinary one-way ANOVA with Dunnett’s multiple comparison test versus Ctrl-EVs was used. \**p* < 0.05; n = 4-7.

These findings indicate that BL-EVs not only display improved uptake but also carry a functionally pro-inflammatory signature. Importantly, the lipidomic remodeling we observed particularly the enrichment of saturated FA and reduced unsaturation may contribute to this phenotype by promoting the formation of rigid lipid microdomains that facilitate receptor clustering and signaling. Such changes could enhance the packaging of pro-inflammatory cargo (e.g., iNOS) into EVs or stabilize EV-cell membrane interactions, thereby amplifying downstream immune activation [64, 73].

### 3.7. Bacterial-stimuli-treated EVs induce cytokine and adhesion molecule expression in endothelial cells

On top of the mediating local immune communication, EVs can disseminate systemically through the bloodstream and lymphatic system [9, 74], where they encounter vascular epithelial cells and may act as modulators of vascular activation.

To explore this, we treated murine endothelial cell line with bacteria-stimulated or Ctrl-EVs for 24 h. We included LPS-EVs as a benchmark, given the strong pro-inflammatory capacity of LPS compared with the more complex BL stimulus. Indeed, LPS-EVs induced a statistically significant increase in IL-6 secretion, whereas BL-EVs did not alter IL-6 levels (Fig. 6d). As discussed above, residual bacterial contaminants should not account for these effects. Because IL-6 acts as a pleiotropic cytokine regulating immune responses, acute inflammation, and chronic disease, its induction in endothelial cells indicates a transition to a pro-inflammatory state that facilitates immune cell adhesion and recruitment. This hypothesis was supported by the significantly elevated production of RANTES (Regulated upon Activation, Normal T-cell Expressed and Secreted; also known as CCL5) after LPS-EV treatment, and a nonsignificant increase following BL-EVs (Fig. 6e). Similarly, ICAM-1 expression was significantly upregulated in endothelial cells after treatment with both LPS- and BL-EVs (Fig. S9). CCL5/RANTES is a chemokine critical for recruiting T cells, monocytes, and macrophages to inflammatory niches, while ICAM-1 serves as an adhesion receptor required for firm leukocyte binding and transmigration across the endothelium. The coordinated induction of RANTES and ICAM-1 indicates an endothelial activation program driven by bacteria-stimulated EVs. We note that BL-EVs induced ICAM-1 but not IL-6, which could be interpreted as a selective endothelial activation profile. This divergence likely reflects different signaling thresholds and kinetics between adhesion molecules and soluble cytokines, an aspect that deserves further study.

Taken together, our finding on MS-1 cells demonstrate that EVs generated under bacterial stimulation, particularly LPS-EVs, are not passive carriers but active drivers of vascular activation, enabling endothelial cells to recruit leukocytes from the systemic circulation to sites of infection. These observations are consistent with prior reports of endothelial activation by EVs and particularly emphasize the stronger effects of EVs from bacteria-infected macrophages [69]. Comparable mechanisms have also been described for EVs derived from *Mycobacterium tuberculosis*–infected macrophages [75], further reinforcing our findings. To our knowledge, this is the first demonstration that BL-EVs can induce endothelial adhesion molecule expression, although weaker than LPS-EVs, indicating that bacterial stimulus type influences the strength of the pro-inflammatory effect.

### 3.8. *In vivo* identification of iNOS-loaded EVs in murine peritoneal disease models and human peritonitis

To validate our *in vitro* observations of iNOS packaging into EVs, we extended our analysis to *in vivo* models of peritoneal inflammation and peritoneal inflammation-associated fibrosis. To further assess the clinical relevance of these findings, we also analyzed EVs isolated from peritoneal exudate samples obtained from patients with acute secondary peritonitis.

In the peritoneal inflammation (PI) mouse model, acute inflammation was induced by intraperitoneal injection of either LPS, a standard inducer of PI and sepsis [26], or BL, the same treatments used for activation of macrophages prior EV isolation in the *in vitro* part of study (Fig. 7a). Both stimuli successfully triggered inflammation, as evidenced by leukocyte redistribution in the peritoneal lavage: lymphocyte counts were significantly reduced, while neutrophils increased robustly and monocytes showed a moderate, non-significant rise (Fig. 7b). LPS further induced systemic inflammation, reflected by altered leukocyte subsets in peripheral blood (Fig. S10). This confirms that our *in vivo* model recapitulates both local and systemic aspects of inflammation.

**Figure 7.**
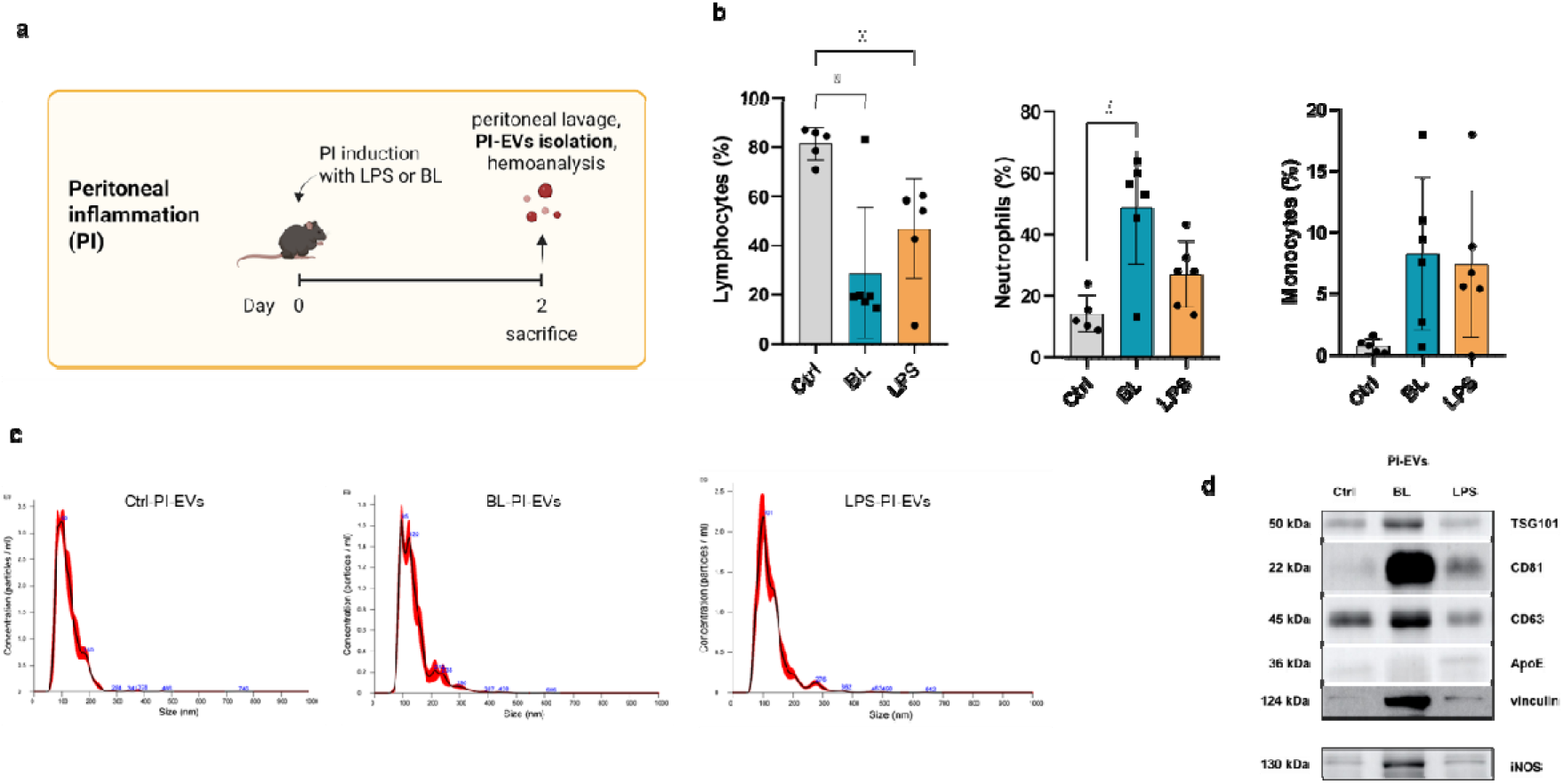
*In vivo* identification of iNOS-loaded EVs in murine peritoneal inflammation (PI) model. **(a) Scheme of *in vivo* experimental setup.** **(b) Leukocyte composition in peritoneal lavage 2 days after induction of PI.** Mice received intraperitoneal injections of lipopolysaccharide (LPS), bacterial lysate (BL), or PBS (Ctrl). Leukocyte subsets were quantified using a clinical hemocytometer and are presented as percentage (mean ± SD). Ordinary One-way ANOVA followed by Bonferroni’s multiple comparisons test versus Ctrl was performed. \**p* < 0.05; n = 5-6. **(c) Particle size and concentration of PI-EVs measured by Nanoparticle Tracking Analysis (NTA).** EVs isolated from peritoneal lavage of mice received intraperitoneal injections of LPS, BL, or PBS (Ctrl). **(d) Detection of EV markers and iNOS in peritoneal inflammation–derived EVs (PI-EVs).** EVs were isolated from lavage fluid of BL-, LPS-, or PBS- (Ctrl) treated mice. Expression of typical EV markers, including negative markers, and iNOS was analyzed by Western blotting. Representative blots are shown.

Given the leukocyte shift characterized by increased neutrophils (and a non-significant rise in monocytes), we hypothesized that inflammatory activation might alter phagocyte-derived EV production and cargo. To test this, we quantified the numbers of EVs isolated from peritoneal lavage (PI-EVs) and found that they did not differ significantly between controls and either LPS- or BL-induced groups. Across all conditions, the PI-EVs displayed a size distribution typical of small EVs (Fig. 7c). Classical EV markers (TSG101, CD63, CD81, vinculin) were detected, together with minor ApoE signal, which we attribute to low-level lipoprotein contamination commonly seen in body-fluid EV preparations (Fig. 7d). Importantly, PI-EVs from both LPS- and BL-treated mice exhibited elevated iNOS. Given that monocytes/macrophages are the main source of iNOS, we infer that the signal originates from macrophage-derived EVs.

Chronic inflammation is connected with various pathologies, including cardiovascular disease, atherosclerosis, and fibrosis. To place our findings on altered EV cargo into a broader context, we selected the peritoneal fibrosis (PF) model as a representative example. In humans, PF is frequently associated with adhesion formation after surgery or intra-abdominal inflammation and is further characterized by tissue hypoxia [76–78]. Using combined hypoxia (CoCl₂) and mechanical stimulation, we induced inflammation and fibrosis in mice, verified by adhesion formation and histology at day 10 (Fig. S11). As early events are crucial for fibrosis initiation [76], we analyzed PF-EVs at day 3 (Fig. 8a). These PF-EVs displayed typical morphology (Cryo-EM) (Fig. 8b) and EV markers (Fig. 8c). Notably, only PF-EVs from fibrosis-induced mice contained elevated iNOS (Fig. 8c). This finding extends our observations from acute inflammation to a clinically relevant chronic pathology, underscoring the robustness of iNOS packaging into EVs.

**Figure 8.**
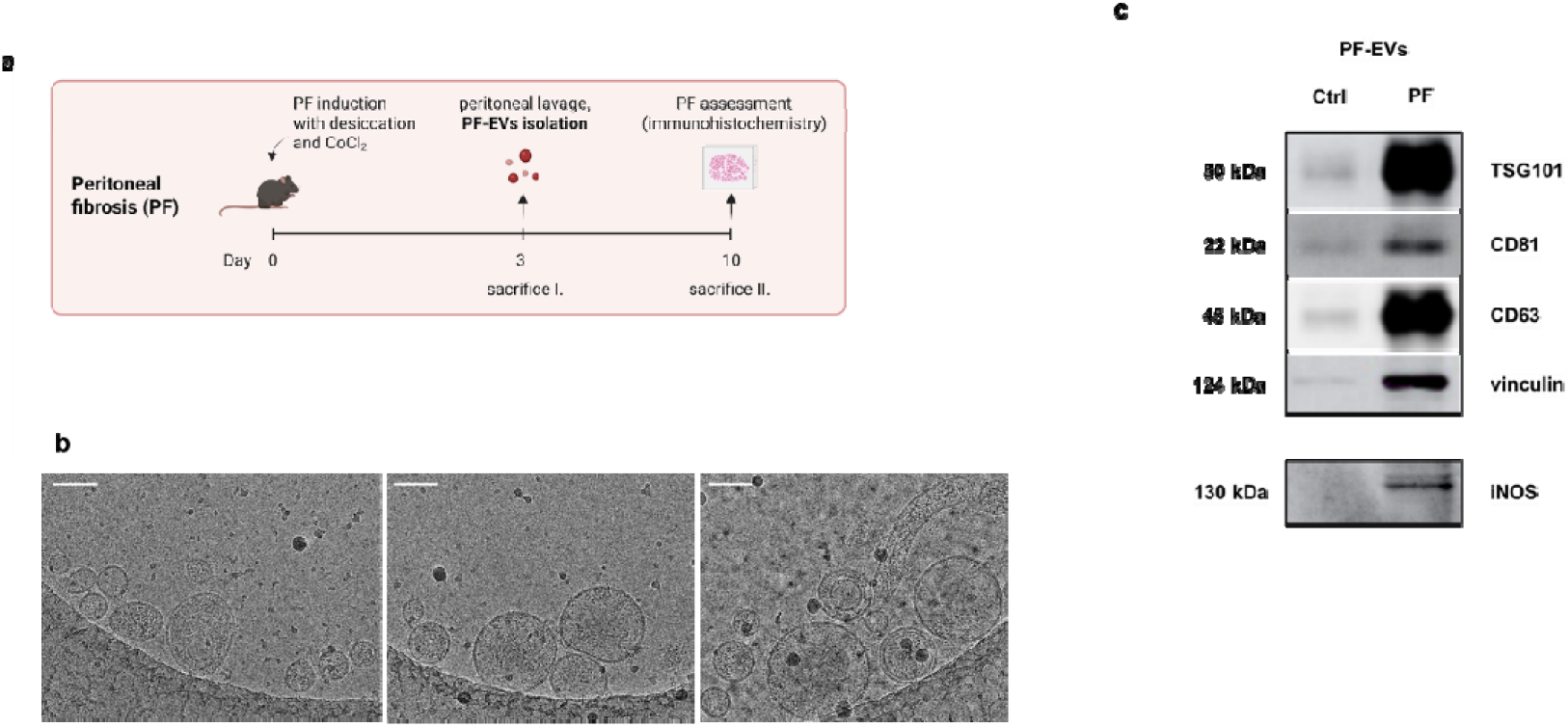
*In vivo* identification of iNOS-loaded EVs in murine peritoneal fibrosis (PF) model. **(a) Scheme of *in vivo* experimental setup.** **(b) Cryo-electron microscopy of PF-EVs.** EVs display characteristic bilayer membrane morphology. Scale bar = 100 nm. **(c) EV markers and iNOS in PF–derived EVs (PF-EVs).** EVs were isolated from the lavage fluid of PF-induced mice and EV markers and iNOS were detected by Western blotting. Representative blots are shown.

To explore the clinical relevance of our findings, we isolated EVs from peritoneal exudates obtained from a pilot patient cohort with surgically confirmed diffuse purulent secondary peritonitis (Fig. 9a). Microbiological culture and NGS confirmed the presence of both Gram-positive and Gram-negative bacterial species in these samples (Table S3). CryoEM and NTA demonstrated that the isolated vesicles exhibited a characteristic lipid bilayer structure (Fig. 9b) and were predominantly within the size range of small EVs (Fig. 9 b, c). Western blot analysis further revealed the presence iNOS associated with patient-derived EVs (Fig. 9d), consistent with our observations in bacteria-stimulated macrophage-derived EVs. To our knowledge, this is the first report demonstrating the *in vivo* association of iNOS transport with small EVs, based on analyses of samples from mouse models (acute peritonitis and peritoneal fibrosis), and human peritoneal exudates from patients with acute peritonitis. Previous work by Webber’s group identified iNOS in circulating microvesicles during sepsis, where they proposed MV-associated iNOS drives excessive NO release and contributes to organ dysfunction [9, 10]. Our study adds two critical advances: (i) we show that iNOS is not limited to larger microvesicles but is also packaged into small EVs, and (ii) we demonstrate this in both acute and fibrosis-associated inflammatory settings.

**Figure 9.**
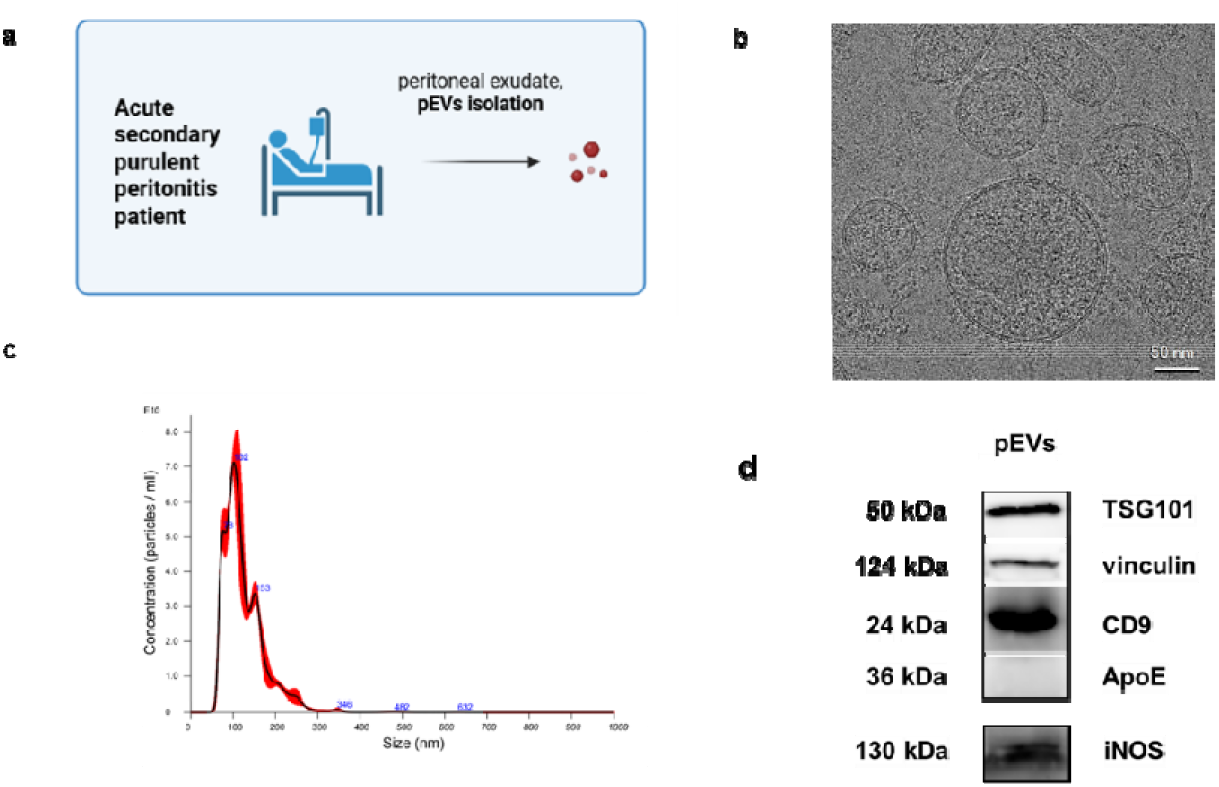
Clinical study cohort of acute secondary peritonitis. **(a) Scheme of experimental setup.** **(b) Cryo-electron microscopy of pEVs**. EVs display characteristic bilayer membrane morphology. **(c) Particle size and concentration of EVs isolated from peritonitis exudate measured by Nanoparticle Tracking Analysis (NTA).** Representative NTA profile of EVs isolated from peritoneal exudates of patients with surgically confirmed acute secondary purulent peritonitis, n=4. **(d) EV markers and iNOS in EVs from acute peritonitis patients (pEVs).** EVs were isolated from peritoneal exudates of patients with surgically confirmed acute secondary purulent peritonitis, and EV markers and iNOS were analyzed by Western blotting. Representative immunoblots are shown, n=4.

## Limitations and Perspectives

This study advances our understanding of how bacterial stimulation remodels macrophage-derived EVs, thereby modulating their uptake and downstream effects, but several limitations should be considered. First, the mechanistic experiments were primarily performed using murine macrophages, endothelial cells, and peritoneal models. Although these systems provide reproducible and mechanistically informative platforms, they may not fully capture human immune or vascular responses. Importantly, the detection of iNOS-positive EVs in EV preparations from peritoneal exudate of patients with peritonitis provides initial clinical relevance; however, larger patient cohorts and functional studies using primary human macrophages and endothelial cells will be required to assess the relevance of these findings in settings such as peritonitis, sepsis, and chronic inflammatory or fibrotic disease. Functionally, although BL-EVs clearly activate macrophages and endothelial cells, it remains uncertain whether these effects depend on EV surface molecules, are mediated by soluble factors within the EV lumen, or result from a combination of both. Clarifying this will deepen our understanding of how EVs propagate inflammatory signals within tissues.

Unfortunately, technical constraints limit our ability to definitively localize EV-associated molecules. Current EV purification methods cannot reliably distinguish luminal cargo from molecules bound to the vesicle surface or co-isolated in non-vesicular structures. Although iNOS is a cytosolic protein and therefore expected to be enclosed within EVs, we have not yet formally demonstrated its intravesicular localization. Despite extensive washing, sucrose cushion separation, and removal of free ligands, we cannot fully exclude low-level contamination with bacterial fragments or non-vesicular aggregates of similar density. Future studies using orthogonal approaches, such as density gradient ultracentrifugation combined with protease protection assays, detergent-mediated vesicle disruption, or immunogold electron microscopy, will be essential to resolve the precise localization of iNOS and other EV-associated molecules.

Looking ahead, although we identify distinct lipid signatures in BL-stimulated EVs, the causal role of these lipids in EV biogenesis, uptake, and inflammatory signaling remains unresolved. Targeted manipulation of lipid pathways, combined with functional assays, will help establish mechanistic links. Ultimately, such studies will clarify how lipid remodeling and iNOS incorporation shape EV-mediated communication and evaluate their potential as biomarkers or therapeutic targets in infection-driven and chronic inflammatory diseases.

## Conclusion

In this study, we demonstrate that bacterial stimulation fundamentally reprograms the molecular composition and biological activity of macrophage-derived small EVs. Bacterial activation drives coordinated remodeling of EV lipid composition, promotes the incorporation of enzymatically active iNOS as a previously unrecognized EV cargo, and enhances the capacity of EVs to propagate inflammatory signaling to macrophages and endothelial cells. The detection of iNOS-positive EVs in murine models of peritoneal inflammation and fibrosis, as well as in EV preparations from patients with acute peritonitis, further supports the physiological and clinical relevance of these findings.

Collectively, our findings establish macrophage-derived EVs as active mediators of bacteria-driven inflammatory communication rather than passive byproducts of macrophage activation. By identifying enzymatically active iNOS as a functional EV cargo and linking its acquisition to coordinated lipid remodeling and enhanced inflammatory activity, this study advances our understanding of how innate immune activation reshapes EV biology. These molecular signatures may serve as accessible indicators of macrophage activation in inflammatory diseases and provide a rationale for exploring EV biogenesis, cargo loading, and EV-mediated intercellular communication as therapeutic targets to limit pathological inflammation.

## List of abbreviations

BL: bacterial lysate from Lacticaseibacillus rhamnosus CCM7091
CFSE: carboxyfluorescein diacetate N-succinimidyl ester
CL: cardiolipin
DGDG: digalactosyldiglyceroles
DB: double bonds
EVs: extracellular vesicles
FA FAs: fatty acid
fPBS: filtered phosphate-buffered saline
iNOS: inducible nitric oxide synthase
LION: Lipid Ontology
LPS: lipopolysaccharide
LTA: lipoteichoic acid
LPCs: lysophosphatidylcholines
MFI: median of fluorescence intensity
MTBE: methyl-tert-butyl ether
NTA: nanoparticle tracking analysis
NO: nitric oxide
LML: number of live maternal cells per mL
PAMPs: pathogen-associated molecular patterns
PRRs: pattern recognition receptors
PF: peritoneal fibrosis
PI: peritoneal inflammation
PE: phosphatidylethanolamine
PGs: phosphatidylglycerols
PCs: phosphatidylcholins
Pis: phosphatidylinositols
RFU: relative fluorescence units
UC: ultracentrifugation
WB: western blot

## Author contributions

Miriam Dutkova: Conceptualization; writing—original draft; writing—review and editing; investigation; data curation; methodology; formal analysis.

Julia Orlovska: Investigation; data curation; methodology.

Kristyna Turkova: Conceptualization; writing—original draft; writing—review and editing; project administration.

Eva Vojtkova: Investigation; data curation; methodology.

Marek Oslacky: Investigation; data curation.

Lucia Calekova: Investigation.

Veronika Mlcekova: Investigation.

Stepan Strnad: Data curation; methodology; formal analysis.

Vladimir Vrkoslav: Data curation; methodology; formal analysis.

Vratislav Berka: Methodology, formal analysis.

Josef Chudacek: Clinics.

Stefan Kolcun: Clinics.

Eva Kriegova: Clinics; data curation.

Lukas Kubala: Supervision; writing—review and editing.

Gabriela Ambrozova: Conceptualization; writing—original draft; project administration; investigation; supervision; methodology; data curation; writing—review and editing.

## Acknowledgements

We thank Anna Kocurkova and Contipro a.s., for their help with peritoneal fibrosis experiments, Ondrej Naar for technical assistance with flowcytometric analysis and Vojtech Novohradsky for access, training, and support with NanoSight NS300 measurements. We are grateful to Vendula Hlavackova Pospichalova and Lenka Markova for providing the flotillin and anti-human iNOS antibodies, respectively.

This output was produced with the support of the OP JAK project “Returns to Research at the Institute of Biophysics of the Czech Academy of Sciences after a Career Break”, Project Reg. No. CZ.02.01.01/00/24_037/0013767; supported by the Ministry of Health of the Czech Republic in cooperation with the Czech Health Research Council within the National Institute CarDia, project No. NW26A-CARDIA.. We acknowledge CF CryoEM, CF CELLIM, Instruct-CZ Centre, supported by MEYS CR (LM2023042) and European Regional Development Fund-Project Innovation of Czech Infrastructure for Integrative Structural Biology (No. CZ.02.01.01/00/23_015/0008175) and MEYS CR (8J24AT032). Miriam Dutkova is a Brno Ph.D. Talent Scholarship holder – funded by the Brno City Municipality, Julia Orlovska supported for excellent diploma thesis (Grant Agency of Masaryk University - Project no. MUNI/C/0132/2025).

## Data availability statement

The datasets generated during the current study have been deposited in Zenodo and will be made publicly available upon acceptance of the manuscript.

