## Supplements for "Bacterial Stimulation Remodels Macrophage Extracellular Vesicle Lipids and Reveals iNOS as an Inflammatory Cargo"

**Content:**

Figures:

S1. Characterization of bacterial lysate (BL) from *Lacticaseibacillus rhamnosus* CCM7091 composition.

S2. Macrophage activation by bacterial lysate (BL) and lipopolysaccharide (LPS).

S3. Flotilin 2 expression in maternal macrophages.

S4. Scheme of preparation of extracellular vesicles (EVs) samples for lipidomic analysis.

S5. Lipidomic profiling of macrophage membranes.

S6. Lipidomic assessment of potential bacterial contaminants.

S7. Lipid Ontology (LION) enrichment analysis of macrophage membranes.

S8. Fatty acid (FA) composition of macrophage membranes.

S9. ICAM expression in endothelial cells

S10. Leukocyte composition in blood 2 days after induction of peritoneal inflammation (PI).

S11. Characterization of pro-fibrotic changes in peritoneal cavity 10 days after peritoneal adhesion induction in C57BL/6 mice.

Table S1: The list of antibodies used for determination of EV markers and pro-inflammatory proteins by Western blot

Table S2: Particle size and concentration of RAW264.7-derived EV samples measured by Nanoparticle Tracking Analysis (NTA).

Table S3: Clinical study cohort of acute secondary peritonitis.

**Figure S1**

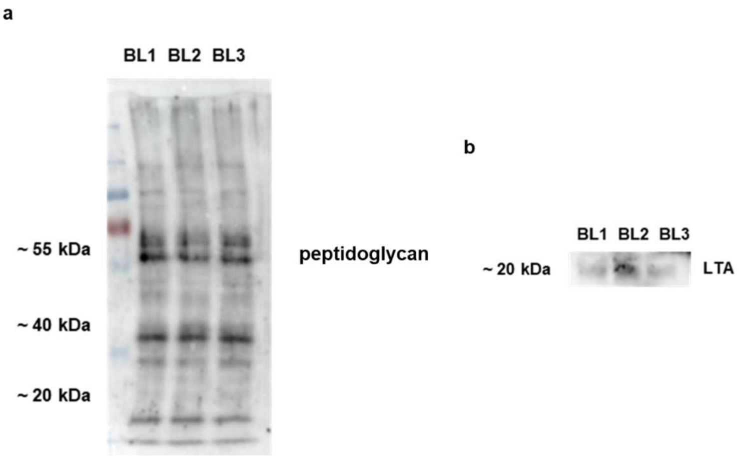


**S1. Characterization of bacterial lysate (BL) from *Lacticaseibacillus rhamnosus* CCM7091 composition. (a)** Peptidoglycan and **(b)** lipoteichoic acid (LTA) were detected in 3 independently prepared BLs by Western blotting. Representative blots are shown.

**Figure S2**

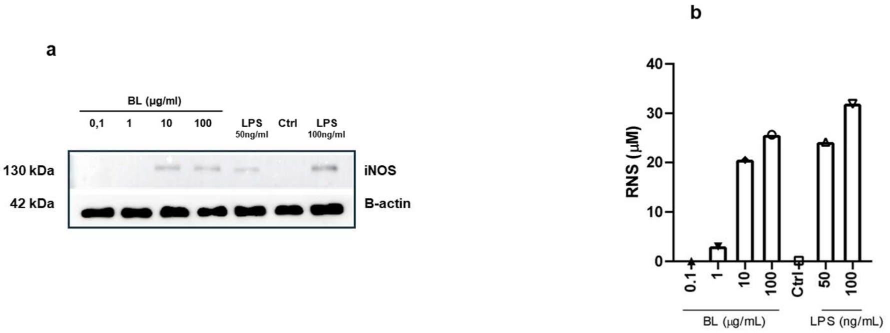


**S2. Macrophage activation by bacterial lysate (BL) and lipopolysaccharide (LPS).** RAW264.7 were treated for 24h with various concentrations of BL (0.1 – 100 ug/mL) or LPS (50 and 100 ng/mL), compared with a negative control (Ctrl, PBS). **(a)** iNOS was detected by Western blotting, together with housekeeping protein (β-actin). Representative blots are shown. **(b)** The levels of reactive species of nitrogen (RNS) were assessed by Griess reaction. One repetition was performed.

#### Figure S3

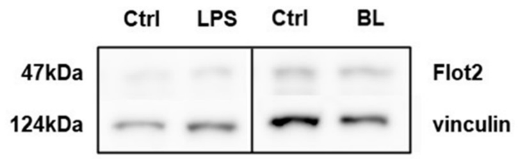


**S3. Flotilin 2 expression in maternal macrophages.** Flotilin 2 and vinculin as a housekeeping protein were detected in RAW264.7 cells treated with lipopolysaccharide (LPS) or bacterial lysate (BL) by Western blotting. One representative repetition is depicted.

#### Figure S4

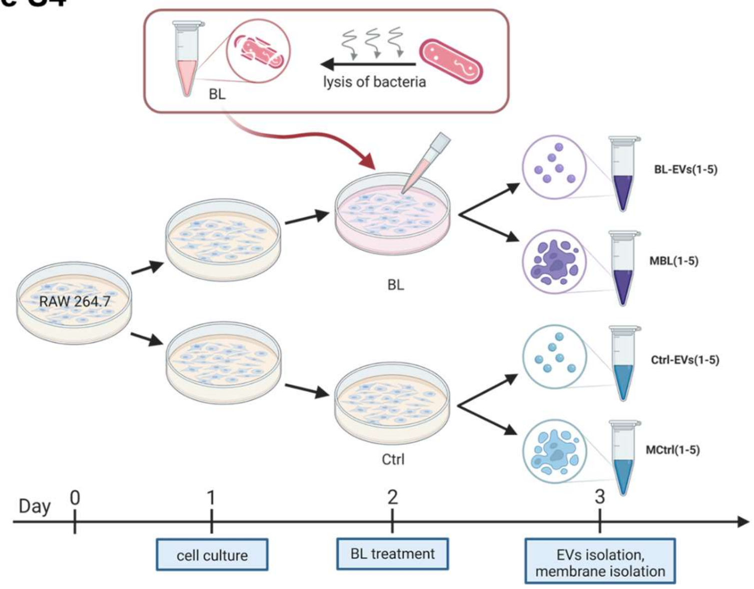


**S4. Scheme of preparation of extracellular vesicles (EVs) samples for lipidomic analysis.** EVs were isolated from bacterial lysate-treated (BL-), or untreated controls (Ctrl-) macrophages. The EVs and maternal macrophage membranes (MBL and MCtrl) were analyzed.

### Figure S5

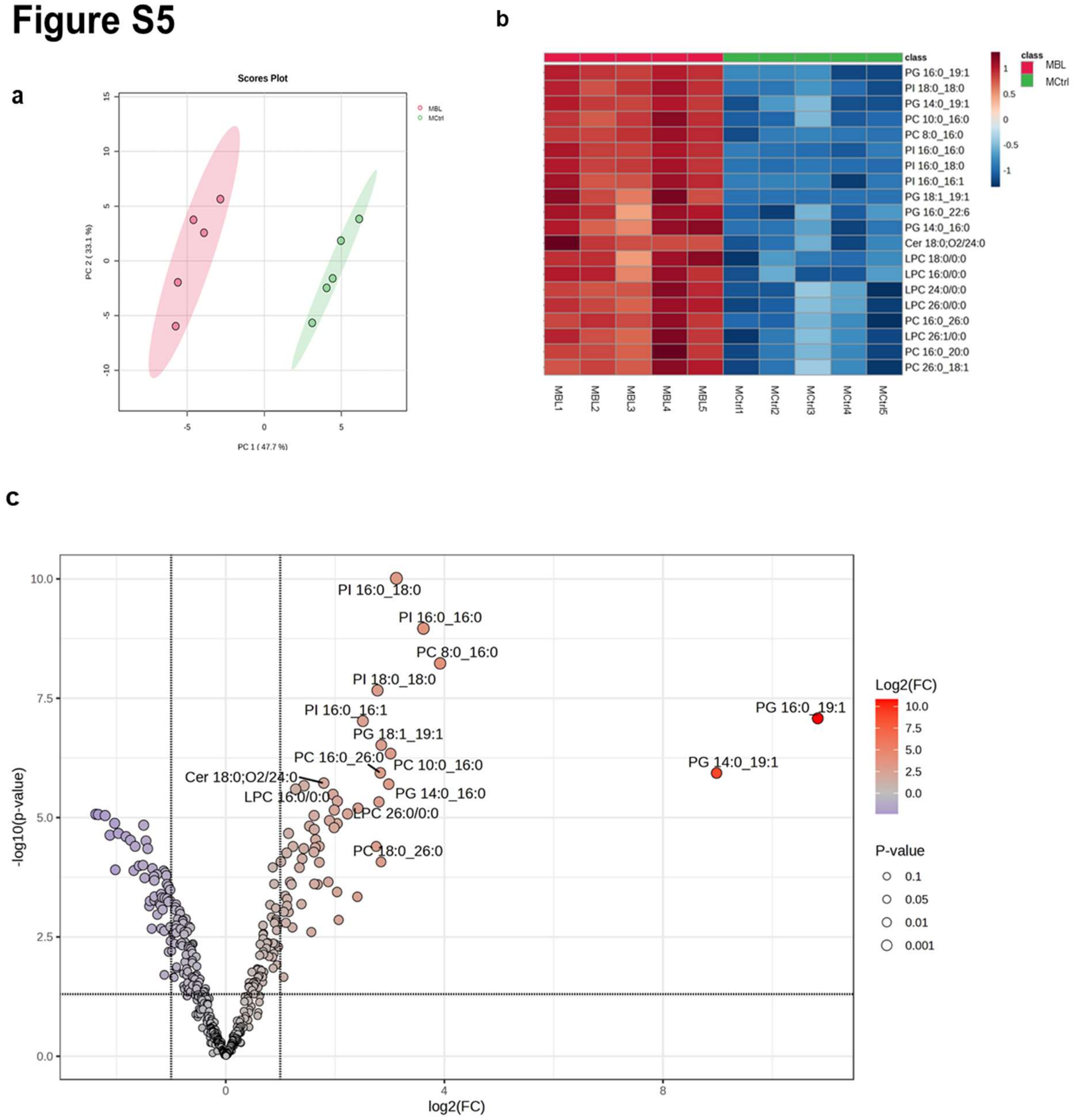


**S5. Lipidomic profiling of macrophage membranes.**

(a) Principal Component Analysis (PCA) of lipid composition in membranes from untreated control (MCtrl) and bacterial lysate-treated (MBL) macrophages.

(b) Heatmap of differentially abundant lipid species in MBL versus MCtrl membranes.

(c) Volcano plot showing significantly altered lipids in MBL versus MCtrl membranes.

**Figure S6**

a

b

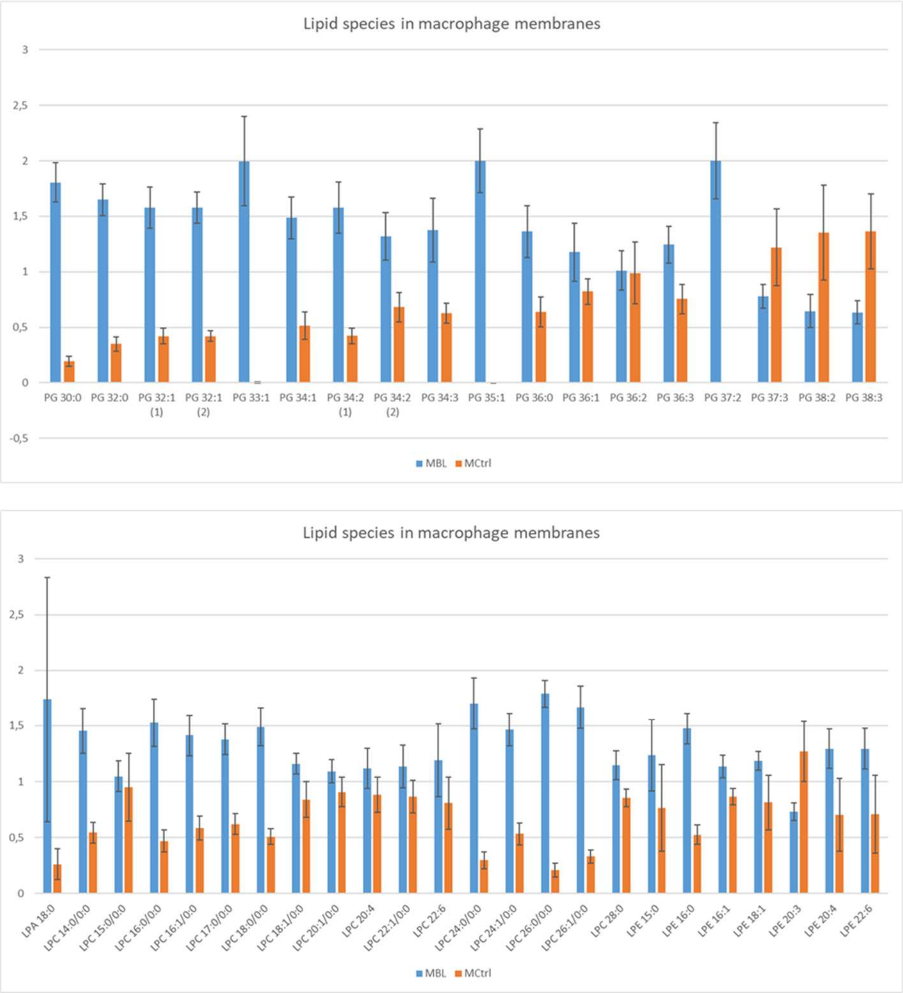


**S6. Lipidomic assessment of potential bacterial contaminants.**

Comparative lipidomic analysis of membranes from untreated control (MCtrl) and bacterial lysate-treated (MBL) macrophages, to evaluate potential bacterial lipid carryover: (a) PGs, (b) lysophospholipids. Data are shown as mean ± SD (n=5).

#### Figure S7

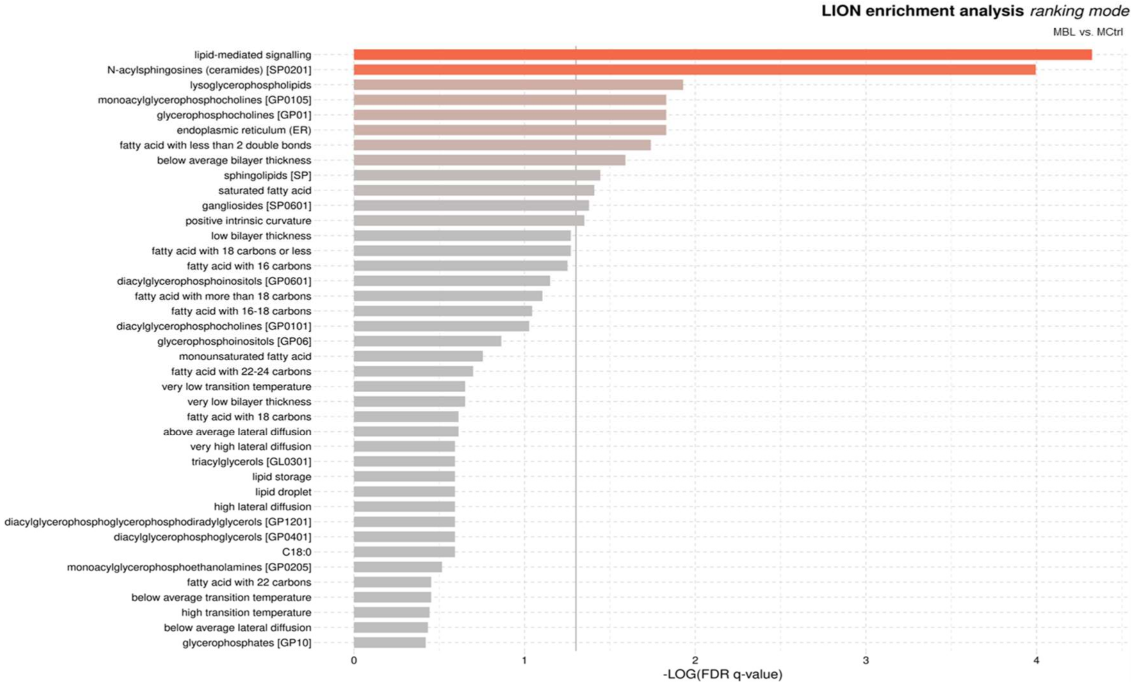


**S7. Lipid Ontology (LION) enrichment analysis of macrophage membranes.**

Functional annotation of lipid classes altered between membranes from untreated control (MCtrl) and bacterial lysate-treated (MBL) macrophages.

#### Figure S8

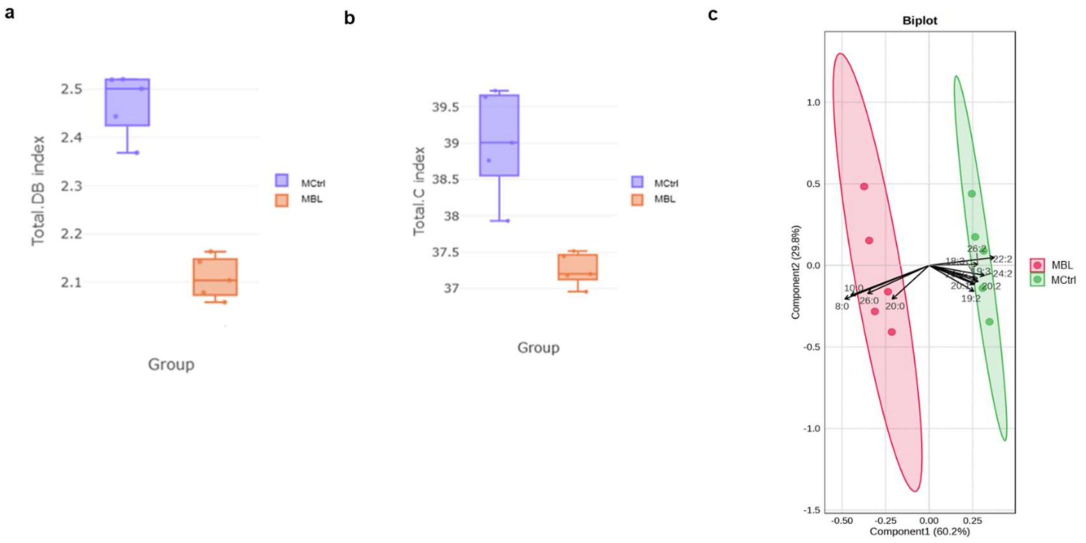


**S8. Fatty acid (FA) composition of macrophage membranes.**

(a) Differences in FA carbon chain length in membranes from untreated controls (MCtrl) and bacterial lysate-treated macrophages (MBL), analyzed using differential expression analysis in LipidSig 2.0.

(b) Differences in FA degree of unsaturation in MCtrl versus MBL membranes, analyzed by LipidSig 2.0.

(c) Biplot of Partial Least Squares Discriminant Analysis (PLS-DA) showing separation of FA composition between MCtrl and MBL samples.

#### Figure S9

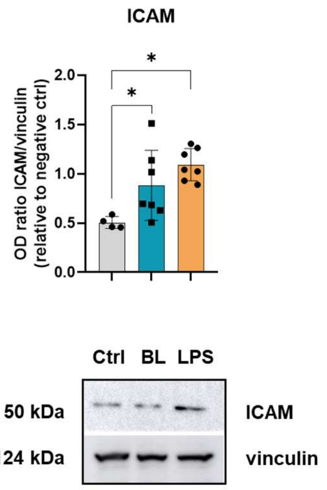


**S9. ICAM expression in endothelial cells.** MS-1 endothelial cell line was treated with Ctrl-, BL-, or LPS-EVs for 24 h. ICAM levels were analyzed by Western blotting. Representative blots for ICAM and vinculin are shown. Densitometry ratios (ICAM/vinculin) were normalized to negative control (vehicle-treated cells, fPBS). Data are presented as mean ± SD. For statistical analysis, Ordinary one-way ANOVA with Dunnett's multiple comparison test versus Ctrl-EVs was used. *$p$ < 0.05; n = 4-7.

#### Figure S10

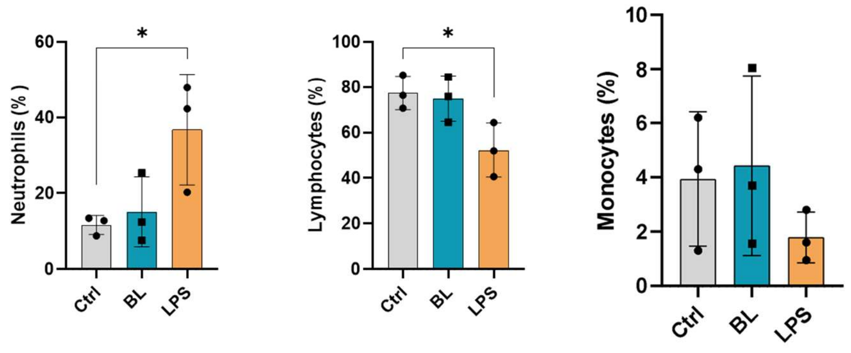


**S10. Leukocyte composition in blood 2 days after induction of peritoneal inflammation (PI).** Mice received intraperitoneal injections of lipopolysaccharide (LPS), *Lacticaseibacillus rhamnosus* CCM7091 bacterial lysate (BL), or PBS (Ctrl). Leukocyte subsets were quantified using a clinical hemocytometer and are presented as percentage (mean ± SD). Ordinary One-way ANOVA followed by Bonferroni's multiple comparisons test versus Ctrl was performed. *$p < 0.05$; n = 3.

#### Figure S11

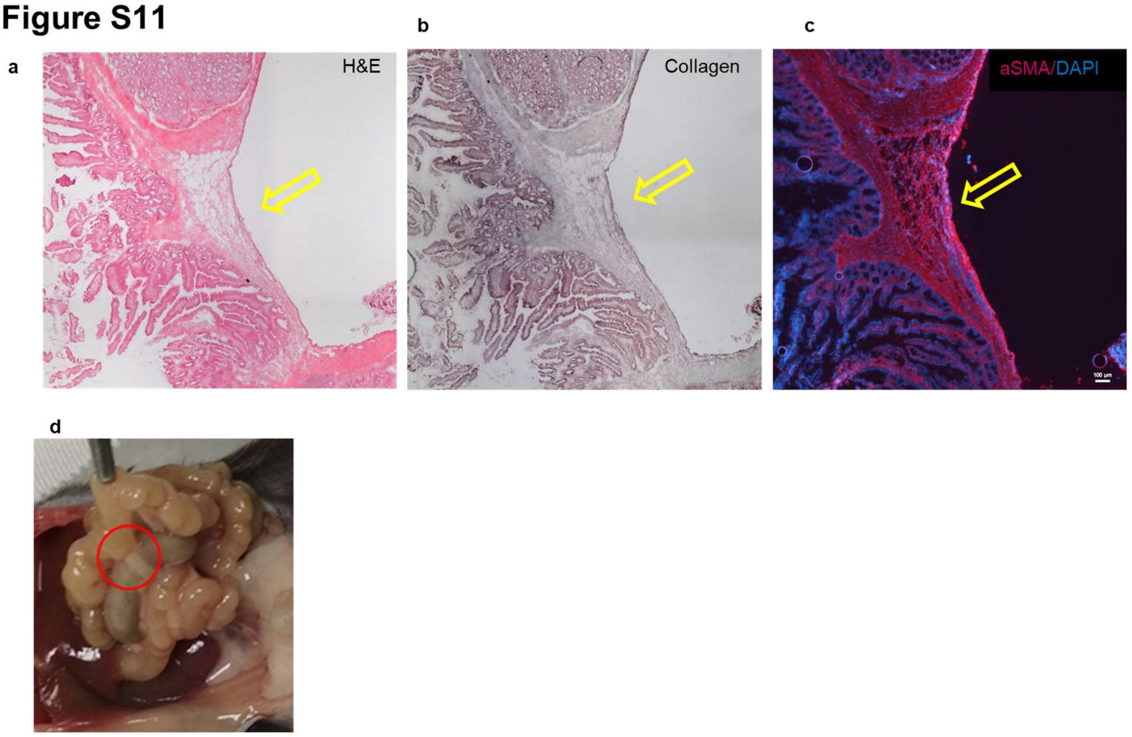


**S11. Characterization of pro-fibrotic changes in peritoneal cavity 10 days after peritoneal adhesion induction in C57BL/6 mice**. (a) Changes in the morphology of tissue determined by hematoxylin and eosin staining, (b) collagen deposition and (c) αSMA expression in newly formed fibrotic tissue. (d) Representative picture of peritoneal adhesion.

**Table S1: The list of antibodies used for determination of EV markers and pro-inflammatory proteins by Western blot**

| Protein | Primary Ab manufacturer | Catalogue number | Blocking solution | Primary Ab dilution | Secondary Ab* dilution | Origin |
|---|---|---|---|---|---|---|
| ***TSG101*** | Santa Cruz Biotechnology, USA | sc-7964 | milk | 1 : 1,000 | 1 : 1,000 | mouse |
| ***CD81*** | Cell Signaling Technology, USA | 10037S | milk | 1 : 500 | 1 : 1,000 | rabbit |
| ***Flotilin 1*** | Abclonal, USA | A3023 | milk | 1 : 600 | 1 : 1,000 | rabbit |
| ***Flotilin 2*** | BD Biosciences, USA | BD 61383 | milk | 1 : 500 | 1 : 1,000 | mouse |
| ***ApoE*** | Santa Cruz Biotechnology, USA | sc-390925 | 5% BSA | 1 : 500 | 1 : 1,000 | mouse |
| ***iNOS*** | BD Biosciences, USA | BD 640431 | milk | 1 : 500 | 1 : 1,000 | mouse |
| ***iNOS + nNOS*** | Abcam, USA | ab202417 | 5% BSA | 1 : 500 | 1 : 1,000 | rabbit |
| ***Vinculin*** | Santa Cruz Biotechnology, USA | sc-139015 | milk | 1 : 1,000 | 1 : 2,000 | rabbit |
| ***LTA*** | Thermo Fisher Scientific, USA | MA1-7402 | milk | 1 : 500 | 1 : 1,000 | mouse |
| ***GAPDH*** | Cell Signaling Technology, USA | 2118 | milk | 1 : 1,000 | 1 : 3,000 | rabbit |
| ***ICAM*** | Santa Cruz Biotechnology, USA | sc8439 | milk | 1 : 1,000 | 1 : 2,000 | mouse |
| ***CD63*** | Santa Cruz Biotechnology, USA | sc-5275 | milk | 1 : 500 | 1 : 1,000 | mouse |
| * all secondary Ab were purchased from Cell Signaling Technology, USA | | | | | | |

**Table S2: Particle size and concentration of RAW264.7-derived EV samples measured by Nanoparticle Tracking Analysis (NTA).** EVs isolated from bacterial lysate-treated (BL-), lipopolysaccharide-treated (LPS-) or untreated control (Ctrl-) macrophages were analyzed.

| | EVs | | | | Maternal cells | | |
|---|---|---|---|---|---|---|---|
| Treatment | Mean particle size (nm) | Mode particle size (nm) | Concentration (particles/mL) | Concentration (particles/frame) | (RAW 264.7) Viability (%) | particles/mL/LML | particles/frame/LML |
| Ctrl-EVs | 146.2 ± 2.4 | 131.7 ± 2.2 | 5.44E+10 ± 1.05E+09 | 129.8 ± 4.5 | 91,7 | 5,93E+10 | 141,5 |
| | 113.3 ± 1.4 | 72.7 ± 8.6 | 2.29E+11 ± 9.78E+09 | 111.6 ± 6.8 | 95,5 | 2,40E+11 | 116,9 |
| | 117.5 ± 1.1 | 76.2 ± 1.0 | 7.66E+10 ± 7.52E+09 | 140.2 ± 13.4 | 95,5 | 8,02E+10 | 146,8 |
| | 130.3 ± 2.5 | 70.6 ± 7.8 | 8.42E+10 ± 5.99E+09 | 177.3 ± 8.5 | 95,2 | 8,85E+10 | 186,3 |
| | 98.9 ± 1.2 | 70.2 ± 8.1 | 9.72E+10 ± 1.26E+10 | 81.3 ± 14.2 | 96,7 | 1,01E+11 | 84,1 |
| | 117.2 ± 3.2 | 70.2 ± 2.6 | 9.05E+10 ± 4.42E+09 | 176.8 ± 2.3 | 97,4 | 9,29E+10 | 181,6 |
| | 116.3 ± 5.3 | 77.2 ± 0.8 | 7.50E+10 ± 2.82E+09 | 150.7 ± 3.8 | 97,5 | 7,69E+10 | 154,6 |
| | 105.1 ± 3.4 | 68.7 ± 13.8 | 8.08E+10 ± 1.16E+09 | 102.0 ± 3.8 | 97,8 | 8,26E+10 | 104,3 |
| | 159.1 ± 3.3 | 72.2 ± 5.7 | 5.39E+10 ± 7.58E+09 | 124.4 ± 16.3 | 95,9 | 5,62E+10 | 129,7 |
| **Average** | **122.7 ± 19.4** | **78.9 ± 20.0** | **9.35E+10 ± 5.29E+10** | **132.7 ± 32.4** | **95.9 ± 1.8** | **9.74E+10 ± 5.53E+10** | **138.4 ± 33.8** |
| BL-EVs | 107.1 ± 0.4 | 73.1 ± 1.8 | 6.14E+10 ± 3.40E+09 | 100.7 ± 7.5 | 97,0 | 6,33E+10 | 103,8 |
| | 123.8 ± 3.7 | 98.8 ± 5.2 | 6.12E+10 ± 4.91E+09 | 41.9 ± 4.6 | 94,9 | 6,45E+10 | 44,2 |
| | 116.0 ± 1.9 | 74.4 ± 1.7 | 7.60E+10 ± 4.45E+09 | 137.9 ± 5.7 | 96,7 | 7,86E+10 | 142,6 |
| | 104.0 ± 2.1 | 71.8 ± 1.3 | 1.09E+11 ± 7.50E+09 | 148.3 ± 15.2 | 97,4 | 1,12E+11 | 152,3 |
| | 105.4 ± 5.6 | 77.6 ± 8.0 | 1.70E+10 ± 3.13E+09 | 20.1 ± 1.0 | 94,1 | 1,81E+10 | 21,4 |
| | 135.1 ± 0.8 | 87.5 ± 2.2 | 6.89E+10 ± 2.68E+09 | 137.9 ± 6.7 | 97,4 | 7,07E+10 | 141,6 |
| | 124.7 ± 1.7 | 94.5 ± 2.6 | 7.20E+10 ± 6.54E+09 | 132.0 ± 12.0 | 97,6 | 7,38E+10 | 135,3 |
| | 103.2 ± 1.8 | 84.7 ± 14.0 | 8.51E+10 ± 1.26E+10 | 60.1 ± 7.4 | 95,2 | 8,94E+10 | 63,2 |
| | 134.6 ± 2.7 | 128.4 ± 17.5 | 7.79E+10 ± 3.84E+09 | 185.6 ± 8.8 | 95,7 | 8,14E+10 | 193,9 |
| | 126.4 ± 0.3 | 83.3 ± 20.6 | 9.28E+10 ± 3.50E+08 | 200.3 ± 2.9 | 97,4 | 9,53E+10 | 205,6 |
| | 117.2 ± 2.9 | 77.1 ± 0.8 | 9.30E+10 ± 1.22E+10 | 182.4 ± 18.5 | 97,6 | 9,53E+10 | 187,0 |
| | 106.3 ± 8.7 | 66.4 ± 4.5 | 1.64E+11 ± 6.36E+10 | 180.2 ± 71.1 | 96,4 | 1,70E+11 | 186,9 |
| | 105.6 ± 2.8 | 75.9 ± 3.8 | 6.06E+10 ± 2.47E+09 | 95.2 ± 10.6 | 97,5 | 6,22E+10 | 97,7 |
| | 97.1 ± 1.3 | 69.5 ± 5.0 | 8.89E+10 ± 3.52E+09 | 86.7 ± 6.3 | 96,1 | 9,25E+10 | 90,2 |
| | 101.8 ± 1.6 | 68.5 ± 2.9 | 7.95E+10 ± 5.17E+09 | 96.2 ± 12.3 | 96,7 | 8,22E+10 | 99,4 |
| **Average** | **113.9 ± 12.4** | **82.1 ± 15.9** | **8.05E+10 ± 3.11E+10** | **120.4 ± 54.9** | **96.5 ± 1.1** | **8.33E+10 ± 3.22E+10** | **124.3 ± 56.4** |
| LPS-EVs | 89.6 ± 1.1 | 77.8 ± 3.0 | 1.80E+10 ± 4.80E+09 | 19.9 ± 0.5 | 95,7 | 1,88E+10 | 20,8 |
| | 118.7 ± 0.3 | 80.6 ± 3.5 | 7.91E+10 ± 2.31E+09 | 151.2 ± 4.3 | 97,5 | 8,11E+10 | 155,1 |
| | 115.3 ± 0.7 | 67.6 ± 1.0 | 1.07E+11 ± 5.31E+09 | 225.1 ± 13.0 | 94,1 | 1,14E+11 | 239,2 |
| | 99.4 ± 1.6 | 70.1 ± 0.9 | 1.22E+11 ± 1.41E+10 | 144.0 ± 12.2 | 96,3 | 1,27E+11 | 149,5 |
| | 122.4 ± 5.8 | 66.5 ± 2.1 | 8.79E+10 ± 5.18E+09 | 189.2 ± 3.7 | 97,1 | 1,26E+11 | 148,3 |
| **Average** | **109.1 ± 14.0** | **72.5 ± 6.3** | **8.28E+10 ± 3.99E+10** | **145.9 ± 77.5** | **96.2 ± 1.3** | **9.32E+10 ± 4.55E+10** | **142.6 ± 78.1** |

**Table S3: Clinical study cohort of acute secondary peritonitis.** Anonymized clinical data included inflammatory markers, comorbidities, infection type, treatment, operative findings and 30-day outcomes.

| Basic characteristics | Patient 1 | Patient 2 | Patient 3 | Patient 4 |
|---|---|---|---|---|
| Age (years) | 25 | 76 | 66 | 56 |
| Sex | Woman | Woman | Woman | Woman |
| Diagnosis | Diffuse purulent peritonitis, and strangulating ileus, 80 cm ileal necrosis | Diffuse purulent peritonitis, perforated appendicitis | Diffuse purulent peritonitis, postoperative day 6 following Miles' procedure | Diffuse purulent peritonitis, fissured rectal carcinoma |
| Ethiology | Lower gastrointestinal tract – ileal necrosis | Lower gastrointestinal tract – perforated appendicitis | Lower gastrointestinal tract | Lower gastrointestinal tract – anal fissure, rectal mass |
| Peritonitis type according to exudate characteristics | Purulent | Purulent | Purulent | Purulent |
| CRP serum levels (mg/l) | 209 | 187 | 249 | 338 |
| Blood Leukocytes (x $10^9$ /l) | 18 | 20 | 12 | 33 |
| Preoperative antibiotics | 0 | 0 | 1 | 0 |
| Microbiology of peritoneal swab (culture + NGS) | *Negative on the day of surgery; Clostridium paraputrificum detected after 24 hours* | *Escherichia coli, Gemella morbillorum, Bacteroides nordii, Enterococcus avium, Parvimonas micra, Porphyromonas gingivalis, Streptococcus constellatus and Bacteroides fragilis.* | *Klebsiella aerogenes, Nakaseomyces glabratus (formerly Candida glabrata), Proteus mirabilis, Staphylococcus epidermidis, Corynebacterium tuberculostearicum* | *Fusobacterium mortiferum, Fusobacterium nucleatum, Parvimonas micra, Peptostreptococcus stomatis, Porphyromonas somerae* |
| ASA score | IIE | IIE | IIIE | IIE |
| qSOFA at admission | 0 | 1 | 1 | 2 |
| SSI (IA, wound) | 1 | 1 | 1 | 0 |
| Sepsis / septic shock | no | Septic shock | Septic shock | no |
| Type of surgical procedure | Vacuum Assisted Closure (VAC/NPWT) | Vacuum Assisted Closure (VAC/NPWT) | Vacuum Assisted Closure (VAC/NPWT) | Vacuum Assisted Closure (VAC/NPWT) |
| Procedure description | Resection of 80 cm of the ileum, side-to-side anastomosis, abdominal cavity lavage, and application of an intra-abdominal VAC | Appendectomy, peritoneal lavage, and application of an intra-abdominal VAC | Surgical re-exploration, peritoneal lavage a negative pressure wound therapy (NPWT) | Rectosigmoid resection (Hartmann's procedure) with end sigmoid colostomy, , and application of an intra-abdominal VAC |
| Length of hospital stay (days) | 6 | 18 | 79 | 29 |
| 90-day mortality | no | no | no | no |
